# Temporal control of inhibition *via* muscarinic acetylcholine receptor signaling generates ON and OFF selectivity in a simple visual circuit

**DOI:** 10.1101/535427

**Authors:** Bo Qin, Tim-Henning Humberg, Anna Kim, Hyong Kim, Jacob Short, Fengqiu Diao, Benjamin H. White, Simon Sprecher, Quan Yuan

**Affiliations:** National Institute of Neurological Disorders and Stroke, National Institutes of Health, Bethesda, MD, 20892, USA; Department of Biology, University of Fribourg, Fribourg, Switzerland; National Institute of Mental Health, National Institutes of Health, Bethesda, MD, 20892, USA

## Abstract

ON and OFF selectivity in visual processing is encoded by parallel pathways that respond to either increments or decrements of light. Despite lacking anatomical features to support split channels, *Drosophila* larvae effectively perform visually-guided behaviors. To understand principles guiding visual computation in this simple circuit, we focus on the physiological properties and behavioral relevance of larval visual interneurons and elucidate their functions in visual processing. We find that the ON vs. OFF discrimination in the larval visual circuit emerges through light-elicited cholinergic signaling that depolarizes the cholinergic interneuron (cha-lOLP) and hyperpolarizes the glutamatergic interneuron (glu-lOLP). Genetic studies further indicate that muscarinic acetylcholine receptor (mAchR)/Gαo signaling in glu-lOLP separates the ON and OFF signals through temporal delays, the disruption of which strongly impacts both physiological responses of downstream projection neurons and dark-induced pausing behavior. Together, our studies identify cellular and molecular substrates for OFF detection in the larval visual circuit and suggests temporal control of inhibition functions as an effective strategy in generating ON and OFF selectivity without anatomical segregation.

## INTRODUCTION

ON and OFF selectivity, the differential neuronal responses elicited by signal increments or decrements, is an essential component of visual computation and a fundamental property of visual systems across species ^1–3^. Extensive studies of adult *Drosophila* optic ganglia and vertebrate retinae suggest that the construction principles of ON and OFF selective pathways are shared among visual systems, albeit with circuit-specific implementations ^4–6^. Anatomically, dedicated neuronal pathways for ON vs. OFF responses are key features in visual circuit construction. Specific synaptic contacts are precisely built and maintained in laminar and columnar structures during development to ensure proper segregation of these signals for parallel processing ^4, 7^. Molecularly, light stimuli elicit opposite responses in ON and OFF pathways through signaling events mediated by differentially expressed neurotransmitter receptors in target neurons postsynaptic to the photoreceptor cells (PRs). This has been clearly demonstrated in the mammalian retina, where light-induced changes in glutamatergic transmission activate ON-bipolar cells via metabotropic mGluR6 signaling and inhibit OFF-bipolar cells through the actions of ionotropic AMPA or kainate receptors ^8, 9^. In the adult *Drosophila* visual system, although both functional and anatomical connectivity have been established for ON and OFF pathways, the molecular machinery mediating signal transductions within these pathways has yet to be identified ^10–13^.

In contrast to the ~6000 PRs and the complex construction of the adult visual system, the much simpler larval *Drosophila* eyes consist of only twelve PRs on each side ^4, 14^. These PRs send axon bundles into the brain and make synaptic connections with a pair of visual local interneurons (VLNs) and approximately ten visual projection neurons (VPNs) in the larval optic neuropil (LON) (Fig. 1a). VPNs further relay signals to higher brain regions that process multiple sensory modalities ^15^. Despite the simple anatomy of their visual systems, larvae excel at visually-guided behaviors. Studies using various paradigms demonstrate that larvae not only rely on vision for negative phototaxis and social clustering, but also form associative memories based on visual cues ^16–21^. How the larval visual system utilizes simple architecture to perform effective information processing is not understood.

**Figure 1.**
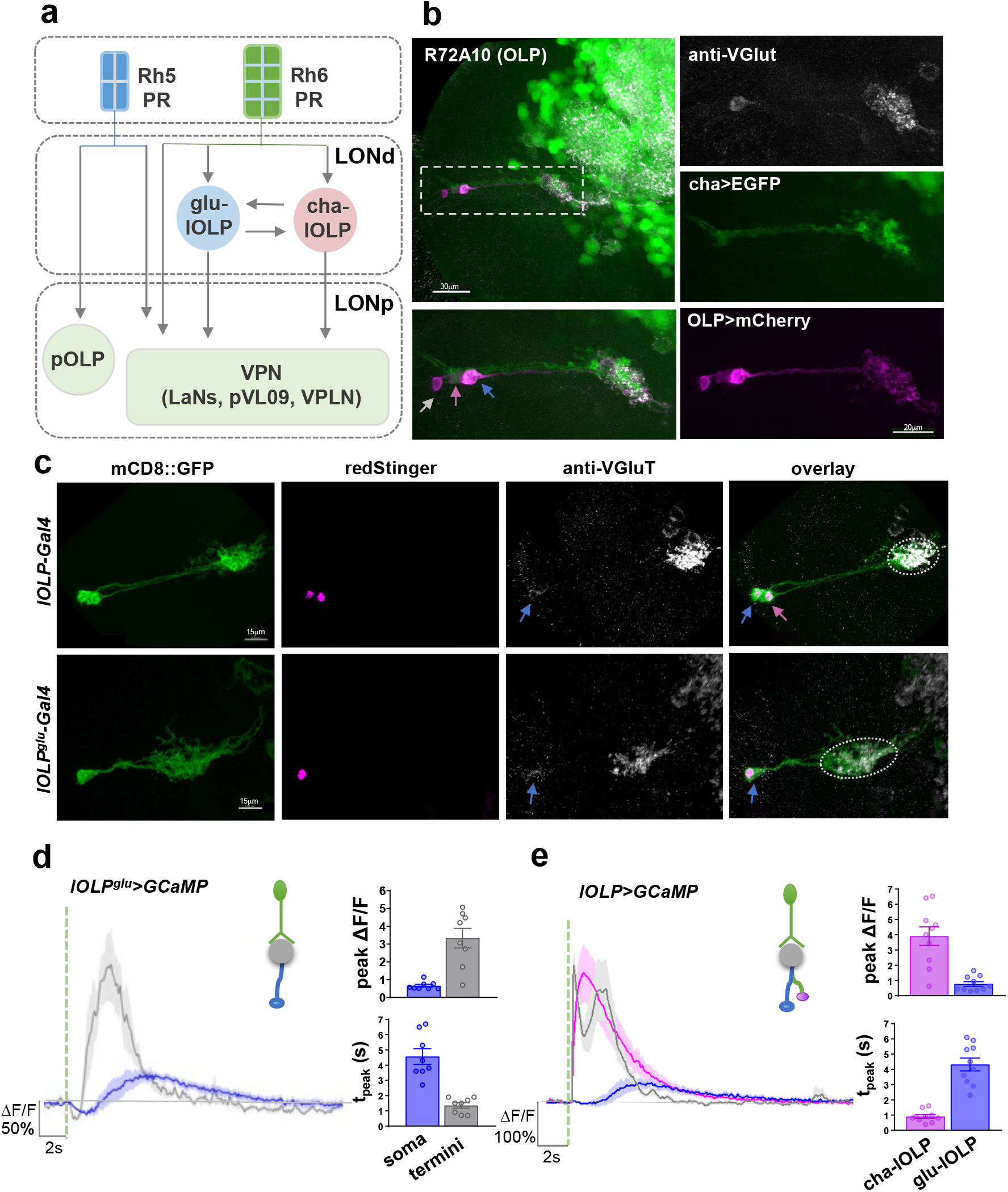
Distinct light-elicited calcium responses in larval visual interneurons. (**a**) Circuit diagram of the *Drosophila* larval visual system. Rh5-expressing photoreceptor neurons (Rh5-PRs) project to the proximal layer of the LON (LONp) and transmit visual signals into the larval brain via direct synaptic connections with visual projection neurons (VPNs). Rh6-PRs project to the distal layer of the LON (LONd) and predominantly form synapses onto two local interneurons, one cholinergic (cha-lOLP) and one glutamatergic (glu-lOLP), which then connect to the VPNs. Gray arrows indicate the unknown effects of light input on OLPs and most VPNs as well as the undetermined interactions between the lOLPs. (**b**) Enhancer screens identified enhancer elements that label three OLPs. R72A10-LexA driven LexAop-mCherry expression (magenta) reveals three somas near the lateral edge of the brain lobe, including the VGluT positive glu-lOLP (blue arrow), the ChAT positive cha-lOLP (pink arrow) and the projection OLP (pOLP, gray arrow). (**c**) Enhancer Gal4 lines specifically labeling two local OLPs (lOLP-Gal4) and the single glu-lOLP (lOLP^glu^-Gal4) were identified. Representative confocal images of larval brains expressing mCD8::GFP and RedStinger driven by enhancer Gal4 lines are shown. Glu-lOLP is positive for anti-VGluT staining in the soma (blue arrows) and terminal processes (dashed circles) that project to the LON. Scale bars = 15μm. (**d-e**) Calcium imaging experiments reveal differential physiological responses to light in two lOLPs. (**d**) Delayed calcium transients in glu-lOLP are observed using GCaMP6f driven by lOLP^glu^-Gal4. The calcium transients obtained at the terminal region (termini) show reduced latency and increased amplitude compared to the ones from the soma. n = 8. (**e**) Light pulses induce fast calcium transients in cha-lOLP (magenta) and slow transients in glu-lOLP (blue). The calcium transient generated at the terminal region is in grey. The average traces of GCaMP6f driven by lOLP-Gal4 and the quantifications of peak value and peak time of changed intensity (ΔF/F) are shown. n = 10. The dashed green line represents a 100 ms light pulse at 561 nm. Shaded areas on traces and error bars on quantifications represent SEM.

A recent connectome study using serial section transmission electron microscopy (ssTEM) reconstruction mapped synaptic interactions within the LON in the 1^st^ instar larval brain ^15^. This comprehensive wiring diagram of the larval visual circuit revealed two separate visual pathways emerging at the level of the PRs that express either blue-tuned Rhodopsin 5 (Rh5-PRs) or green-tuned Rhodopsin 6 (Rh6-PRs). Rh5-PRs project to the proximal layer of the LON (LONp) and form direct synaptic connections with all VPNs, whereas Rh6-PRs project to the distal layer of the LON (LONd) and predominantly target two local interneurons, one cholinergic (cha-lOLP) and one glutamatergic (glu-lOLP). The two PR pathways then converge at the level of VPNs (Fig. 1a). Behavioral studies suggest that the spectral differentiation in PRs does not code for color discrimination, but rather generates independent processing for spatial and temporal information during larval navigation. Specifically, while Rh5-PRs are essential for navigation based on spatial cues, Rh6-PRs are involved in perceiving temporal changes of light intensity during larval head casts and contribute to effective light avoidance behaviors ^15, 17, 21^. Thus, both behavioral and connectome studies indicate the structural and functional segregation of Rh5- and Rh6-PR pathways.

The connectome study also revealed potential functions for cha- and glu-lOLP, the two local interneurons. The lOLPs, together with one of the VPNs, the pOLP, are known as optic lobe pioneer neurons (OLPs), because they are the earliest differentiated neurons in the larval optic lobe ^22–24^. Besides relaying visual information from Rh6-PRs to downstream VPNs, the lOLPs also form synaptic connections with each other and receive neuromodulatory inputs from serotonergic and octopaminergic neurons, suggesting the possibility that lOLPs function as a hub that processes light information and generates feature-specific presentations in downstream VPNs (Fig. 1a). These connectivity patterns led to a proposal that the lOLPs serve as ON and OFF detectors in the larval visual circuit^15^. However, the lack of evidence for anatomical separation at either the input or output level of the lOLPs makes it unclear how they could support differential coding for ON and OFF signals.

Here, building upon these recent findings in anatomical connectivity and behavioral output of the larval visual circuit, we sought to investigate the physiological properties of the pair of lOLPs and determine the molecular machinery underlying their information processing abilities. We identified enhancer lines for targeted expression of genetically-encoded voltage and calcium indicators and observed differential physiological responses towards light increments or decrements in cha-lOLP and glu-lOLP, indicating their functions in mediating ON and OFF discrimination. Furthermore, we found that light-induced inhibition on glu-lOLP is mediated by mAchR-B/Gαo signaling, which not only generates the sign-inversion required for an OFF response, but also encodes temporal information between the cholinergic and glutamatergic transmissions received by downstream VPNs. Lastly, genetic manipulations of glu-lOLP strongly modified physiological responses of VPNs and eliminated dark-induced pausing behaviors. Together, our studies identify specific cellular and molecular pathways that mediates the OFF detection in *Drosophila* larvae and provide evidence for ON and OFF selectivity generated through the temporal control of inhibition, a strategy likely applicable to other circuits constructed without anatomical segregation.

## RESULTS

### Identification and characterization of enhancer Gal4 lines for visual interneurons

To perform physiological and genetic studies on the lOLPs, we first searched for specific Gal4 enhancers. OLP somas are located on the lateral edge of the brain lobe where the optic nerve enters the brain ^22–24^ (Fig. 1b). Their processes extend along the optic nerve and densely innervate the LON. This anatomical feature enabled us to visually screen the enhancer Gal4 collection produced by the Janelia Farm FlyLight Project and identify candidate driver lines labeling OLPs ^25, 26^.

We selected three Gal4 enhancer lines, R72E03, R84E12 and R72A10, based on their differential coverage of OLPs (Fig. 1a, b, Fig. S1). The number and identity of the OLPs labeled by these enhancer lines were determined by staining with anti-ChAT and anti-VGluT antibodies (Fig. 1b, c, Fig. S2) ^22–24^. R72E03-Gal4 (lOLP^glu^-Gal4) labels glu-lOLP only, R84E12-Gal4 (lOLP-Gal4) labels cha- and glu-lOLP, and R72A10-Gal4 (OLP-Gal4) labels both lOLPs and the pOLP. We also tested a R72A10-LexA line which showed the same expression pattern as the Gal4 (Fig. 1b, Fig. S1) ^15^. Single cell labeling using the FLP-out technique and the R84E12-Gal4 enhancer indicate that two lOLPs have similar projection patterns and their termini are largely contained within the LON region (Fig. S2).

### Light stimulation elicits differential calcium responses in OLPs

Having identified enhancer lines that specifically label subsets of OLPs, we then examined their physiological properties with optical recordings. Since OLPs are direct synaptic targets of PRs, we expected to observe light-evoked calcium responses in these neurons. Our previous studies established a protocol to record light-elicited physiological responses in the target neurons of PRs using larval eye-brain explants, in which the Bolwig’s organ, the larval light sensing organ that contains the PRs, the optic nerve and the brain lobe are kept intact ^27^. This approach allows us to deliver temporally controlled light simulations using either the 488 or 561 nm laser while detecting calcium transients with cell-specific expression of GCaMP6f through two-photon imaging (Fig. S3a) ^28^. In addition, to test for the compartmentalization of light-evoked calcium responses, we recorded GCaMP signals in both soma and terminal regions (Fig. 1d) ^29^.

Calcium imaging using lOLP^glu^-Gal4 and lOLP-Gal4 revealed distinct light-elicited physiological responses in the two local OLPs. Upon light stimulation, a 100 ms light pulse delivered by the 561 nm laser, glu-lOLP exhibited a small reduction in the GCaMP signal followed by a slow calcium rise, whereas cha-lOLP responded to light with a large and immediate calcium rise (Fig. 1d, e). Calcium transients obtained from the terminal region of glu-lOLP displayed the same biphasic waveform as those in the somas, but with higher amplitudes and shorter latencies (Fig. 1d). In addition, termini recordings of both lOLPs produced calcium transients with two distinct peaks that clearly reflect the temporal differences in the light-induced responses of the two lOLPs (Fig. 1e). Using R72A10-LexA enhancer-driven LexAop-GCaMP6f expression, we also obtained comparable results for the two lOLPs and characterized the profile of the light response in pOLP, which showed the same initial reduction followed by a slow calcium rise similar to, but with a greater amplitude than, the glu-lOLP response (Fig. 2a, c, d, Fig. S3a, S4). Lastly, we quantified glu-lOLP data sets generated by three different enhancer lines and obtained similar results that indicated the consistency of the results (Fig. S4).

**Figure 2.**
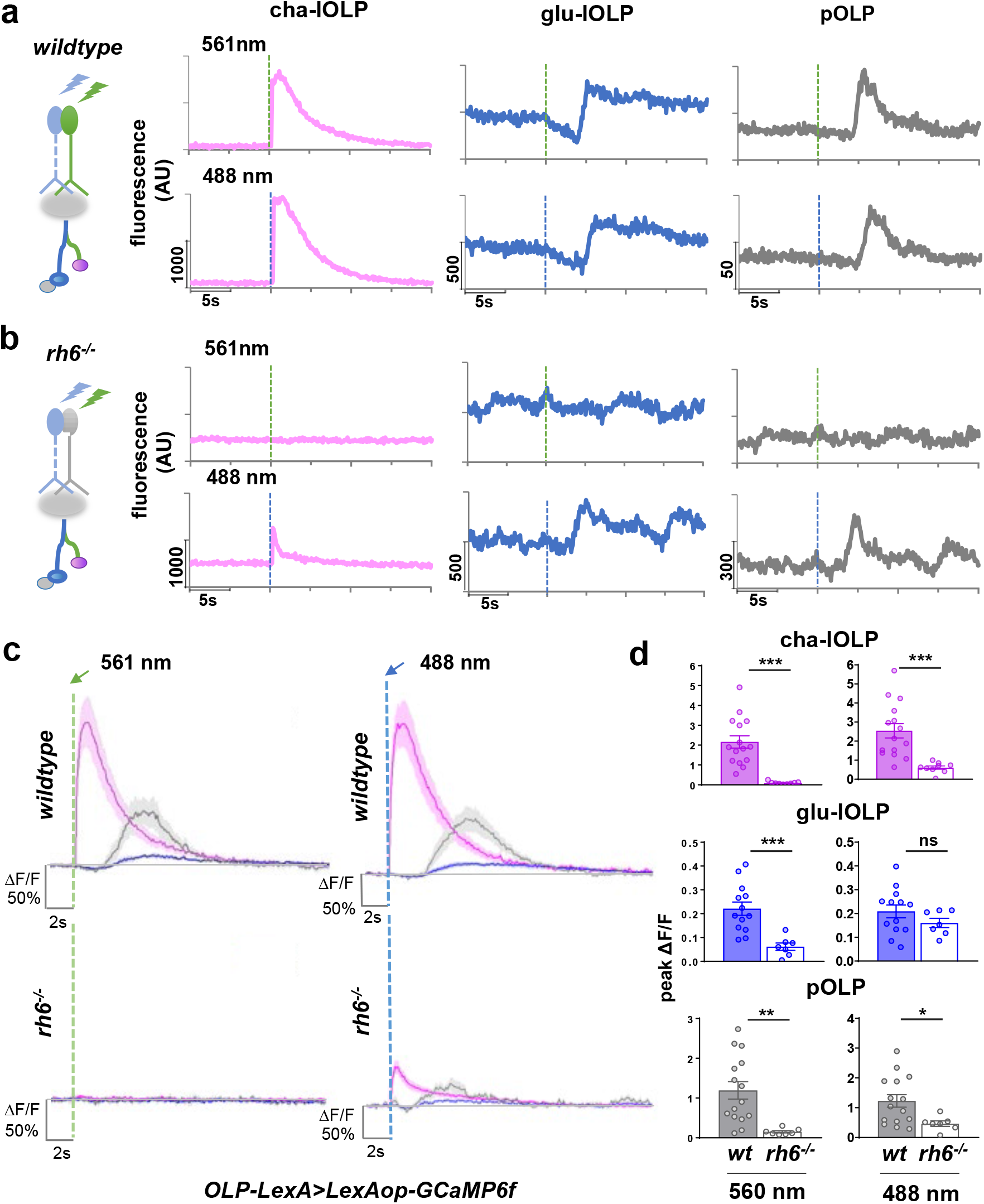
OLPs receive presynaptic inputs predominantly from Rh6-PRs. (**a-b**) The contribution of Rh5- and Rh6-PRs to light-evoked calcium responses in OLPs as revealed by stimulation at different wavelengths in wildtype and Rh6 mutants. Left: schematic diagram illustrating the stimulation scheme used in calcium imaging experiments. Green or blue light pulses (dashed lines, green: 561 nm, blue: 488 nm) activate Rh5- or Rh6-PRs and elicit GCaMP6f signals driven by OLP-LexA in the somas of OLPs. Right: Representative raw traces of OLP>GCaMP6f collected from wildtype and Rh6 mutants (*rh6*^−/−^). Magenta: cha-lOLP, blue: glu-lOLP, grey: pOLP. (**c-d**) OLPs are functionally connected to Rh6-PRs in the 3^rd^ instar larval brain. Light pulses (dashed lines, green: 561 nm, blue: 488 nm) induced fast calcium transients in cha-lOLP (magenta) and slow transients in glu-lOLP (blue) and pOLP (grey). Compared to the wild type controls, OLPs in Rh6 mutants showed no response towards green light (561 nm) stimulation and dampened responses toward blue light (488 nm) stimulation except for glu-lOLP, which remained equally responsive. The **(c)** average traces and **(d)** quantification of peak value of changed intensity (ΔF/F) are shown. Shaded areas on traces and error bars on quantifications represent SEM. Wild type control: cha-lOLP, n = 15; glu-lOLP, n = 13; pOLP, n = 15. Rh6 mutant (rh6^−/−^): cha-lOLP, n = 9; glu-lOLP, n = 7; pOLP, n = 7. cha-OLP, 561 nm: p < 0.0001, t = 5.102, df = 22; cha-OLP, 488 nm: p = 0.0007, t = 3.929, df = 22; glu-OLP, 561 nm: p = 0.0009, t = 3.977, df = 18; glu-OLP, 488 nm: p = 0.2362, t = 1.225, df = 18; pOLP, 561 nm: p = 0.0044, t = 3.207, df = 20; pOLP, 488 nm: p = 0.0261, t = 2.402, df = 20. Statistical significance determined by student’s t-test. *P* ≥ 0.05 was considered not significant, \*\*\**P* < 0.001. \*\**P* < 0.01. \**P* < 0.05.

Together, our calcium imaging studies on the OLPs confirmed their physiological roles as visual interneurons and reveal distinct profiles in their light-evoked responses. Notably, the calcium transients obtained from cha- and glu-lOLP resemble the ones observed in adult fly visual interneurons that belong to either the ON or OFF pathways, respectively, suggesting potential functional similarities between lOLPs and the interneurons in adult visual ganglia ^12, 29^.

### OLPs receive presynaptic inputs predominantly from Rh6-PRs

The ssTEM connectome studies indicate that, in the 1^st^ instar larval brain, the majority of lOLPs’ PR inputs come from Rh6-PRs, while pOLP receives PR inputs directly from Rh5-PRs ^15^ (Fig. 1a). This biased synaptic connection suggests that the OLPs’ light responses might exhibit spectral selectivity. To establish the functional connectivity between subtypes of PRs and OLPs, we performed calcium imaging experiments with light stimulations delivered at 488 nm (blue) and 561 nm (green) wavelengths (Fig. S3a). Previous studies indicated that Rh6 detects light within the 400-600 nm range and its maximal spectral sensitivity is ~437 nm, while Rh5 detects light from 350-500 nm and its maximal spectral sensitivity is ~508 nm. Therefore, Rh6-PRs are sensitive to light stimulations at both 488 nm and 561 nm, whereas Rh5-PRs only respond to blue light at 488 nm ^30^. These features, in combination with a loss-of-function Rh6 mutant (*rh6*^−/−^) ^31^, allowed us to examine the specific contributions of Rh5- and Rh6-PRs to the OLPs’ light responses.

In wildtype larvae, 488 and 561 nm light stimulations elicit almost identical responses from the OLPs (Fig. 2a, c, d). In contrast, all responses to green light (561 nm) were eliminated in the Rh6 mutant (*rh6*^−/−^), demonstrating that green light-evoked responses in OLPs are solely generated by visual transduction in Rh6-PRs (Fig. 2b-d). To test the functional connectivity between Rh5-PRs and OLPs, we performed experiments using blue light (488nm) stimulations in the Rh6 mutant background, where blue light-elicited responses are exclusively generated by Rh5-PRs. Interestingly, compared to the wildtype control, blue light-induced calcium responses in cha-lOLP and pOLP were significantly reduced in Rh6 mutants, suggesting that Rh6-PR is the primary mediator of light inputs into these two neurons. On the other hand, the Rh6 mutation produced no significant difference in glu-lOLP’s blue light response (Fig. 2b-d), indicating that glu-lOLP receives both Rh5- and Rh6-PR inputs.

Together, our findings demonstrate that cha-lOLP and pOLP receive most of their light inputs from Rh6-PRs, while Rh5-PRs provide a minority of functional inputs. In contrast, glu-lOLP has strong functional connections to both Rh5- and Rh6-PRs. These results generated by calcium imaging at the 3^rd^ instar larval stage largely agree with the connectivity map produced in the 1^st^ instar larval brain ^15^, suggesting that biased Rh6-PR/lOLP synaptic connectivity is mostly preserved during larval development and can be detected through functional analyses. On the other hand, the calcium imaging experiments also identified functional connections that were not indicated in the connectome study. Specifically, our results showed that glu-lOLP receives functional inputs from both Rh5- and Rh6-PRs and that pOLP is mainly driven by Rh6-PR input. These differences may be attributed to either developmental changes in circuit connectivity or physiological interactions among neurons that are not directly reflected by anatomical connections, highlighting the importance of complementing connectome analyses using physiological studies.

Additionally, this set of experiments validated our imaging protocols and indicated that, at the intensities we tested, both blue (488 nm) and green (561 nm) light stimulations elicit robust calcium responses in the OLPs (Fig. 2a, c, d). Therefore, light stimulations delivered at either wavelength can be used to study light-evoked responses in the OLPs and downstream VPNs.

### Light induces hyperpolarization in glu-lOLP and depolarization in cha-OLP

GCaMP recording experiments on glu-lOLP revealed an initial reduction followed by a slow rise in calcium signal, suggesting a light-induced suppression of its neuronal activity. To measure light-induced calcium and voltage responses within the same lOLP neurons, we used genetically-encoded voltage sensor Arclight together with the red calcium indicator RCaMP ^32, 33^. By matching calcium profiles with voltage changes, we found that light pulses induce depolarization and fast calcium transients in cha-lOLP but hyperpolarization and slow calcium transients in glu-lOLP (Fig. 3a, b). We compared the RCaMP recordings with the previous GCaMP recordings and found that although the amplitude of calcium transients reported by RCaMP recordings was reduced compared to the results obtained by GCaMP recordings, the waveforms of lOLPs’ calcium responses remain the same (Fig. 3b, Fig. S8).

**Figure 3.**
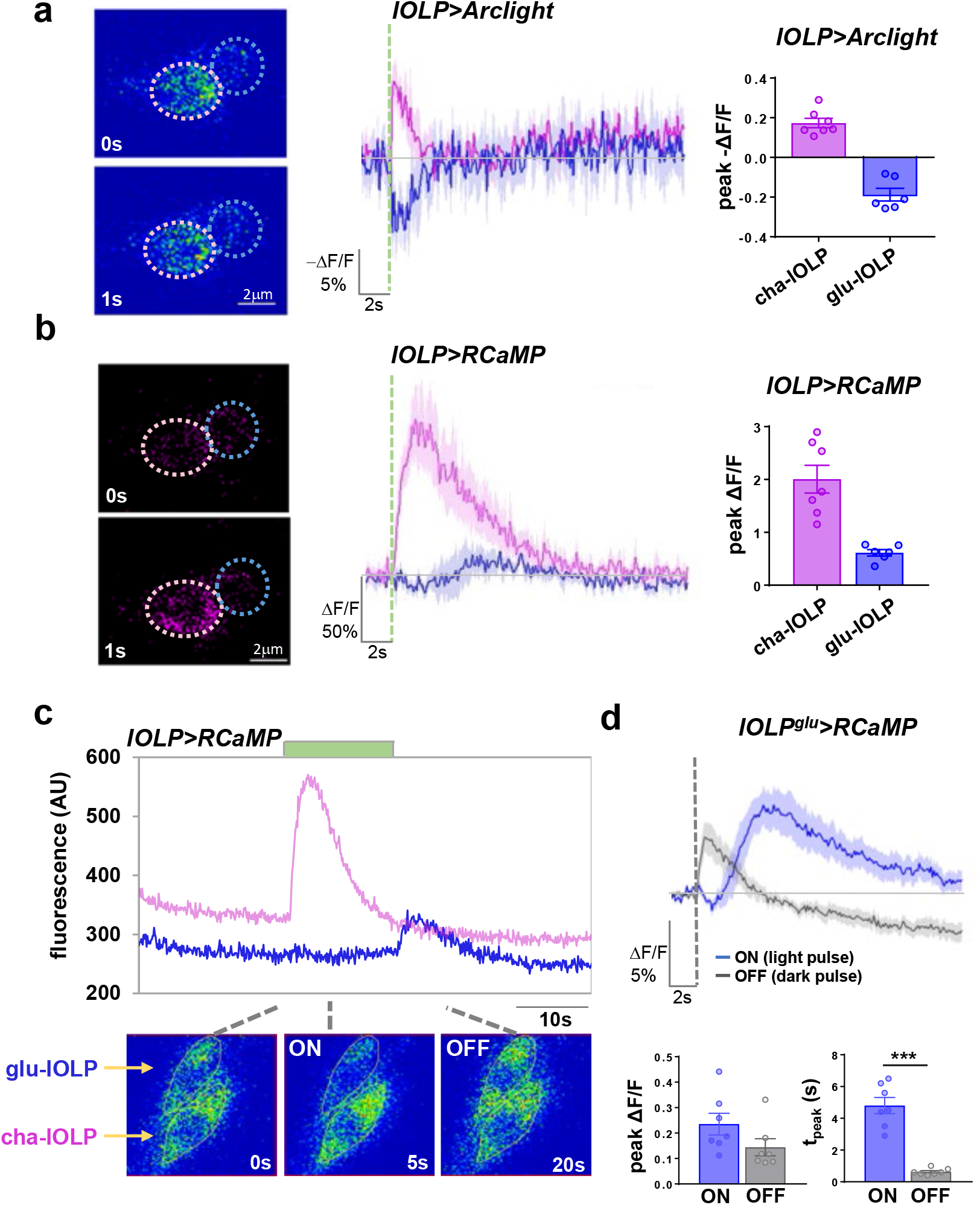
Light activates cha-lOLP and inhibits glu-lOLP. (**a-b**) Optical recordings using the voltage sensor Arclight together with the calcium sensor RCaMP reveal light-induced depolarization and fast calcium transients in cha-lOLP (magenta) as well as hyperpolarization and delayed calcium transients in glu-lOLP (blue). Representative frames from the recordings (left), averaged traces (middle) and the quantification of peak values of the changed intensity (ΔF/F) (right) are shown. Scale bars and time are as indicated. Somatic regions used for quantification are marked by dashed circles. The dashed green line represents a 100 ms light pulse. cha-lOLP, n = 7; glu-lOLP, n = 6. (**c**) cha-lOLP exhibits ON responses while glu-lOLP exhibits OFF responses. A representative raw trace from the lOLP>RCaMP recording is shown (top). The sample was subjected to an extended (12.5s) light stimulation (green bar). cha-lOLP responded to the light onset but showed no response to the light offset. In contrast, the light onset induced a small reduction of calcium signal in glu-lOLP, while the light offset produced an immediate calcium rise. Representative frames of the recording are shown (bottom). (**d**) ON and OFF signals generate calcium transients with different temporal profiles in glu-lOLP. Average traces of calcium transients generated by recordings of lOLP^glu^-Gal4 driving RCaMP are shown, demonstrating the slow calcium response to the light pulse (ON response, blue) and the fast calcium response to the dark pulse (OFF response, grey). The response amplitudes were not significantly different. The average traces (top) and the quantification of peak value and peak time of changed intensity (ΔF/F) (bottom) are as shown. n = 7 in both groups. ON: p = 0.1205; OFF: p < 0.001. Shaded areas on traces and error bars on quantifications represent SEM. The dashed line represents a 100 ms light or dark pulse. Statistical significance determined by student’s t-test. *P* ≥ 0.05 was considered not significant, \*\*\**P* < 0.001.

### Light induces hyperpolarization in glu-lOLP and depolarization in cha-OLP

GCaMP recording experiments on glu-lOLP’s light responses revealed an initial reduction followed by a slow rise in the calcium signal, suggesting a light-induced suppression of its neuronal activity. To measure light-induced calcium and voltage responses within the same lOLP neurons, we examined the change in membrane potential using the genetically-encoded voltage sensor Arclight while recording the calcium transient with the red calcium indicator RCaMP ^32, 33^. By matching calcium profiles with voltage changes, we found that light pulses induce depolarization and fast calcium transients in cha-lOLP but hyperpolarization and slow calcium transients in glu-lOLP (Fig. 3a, b). We compared the RCaMP recordings with the previous GCaMP recordings and found that although the amplitude of calcium transients reported by RCaMP recordings was reduced compared to the results obtained by GCaMP recordings, the waveforms of lOLPs’ calcium responses remain the same (Fig. 3b, Fig. S8).

We next tested how the lOLPs respond to light increments and decrements by monitoring calcium responses during onsets and offsets of extended light exposures. Although the two-photon recordings of GCaMP6f provide the best image quality, it is incompatible with the extended light exposure due to the light sensitivity of the detector. Therefore, in the following experiments, we used RCaMP as the calcium indicator, which can be imaged using a confocal laser tuned to 561 nm with low intensity, reducing the effects of photobleaching on both the calcium sensor and the photoreceptors. Additionally, this protocol allowed us to alter the light cycles or deliver a dark pulse by tuning the 488 nm laser during imaging sessions (Fig. S5a).

RCaMP recordings showed that, with an extended light exposure, cha-lOLP only responded to the light onset with a fast calcium transient, demonstrating its specific response to light increments. In contrast, the light onset induced a small yet noticeable reduction of calcium signal in glu-lOLP, while the light offset produced an immediate calcium rise, suggesting that glu-lOLP is activated by the light decrements (Fig. 3c). This observation is consistent with light hyperpolarizing glu-lOLP at the onset of the light exposure and generating a sustained inhibition until the offset of the light exposure.

We performed additional experiments to examine the differential responses of glu-lOLP toward light increments and decrements by subjecting the preparation to 100 ms light or dark pulses that are contrast matched. We found that a 100 ms dark pulse, or a brief reduction in light intensity following an extended light exposure, is sufficient to generate an immediate calcium rise in glu-OLPs. Compared to the slow calcium responses induced by 100 ms light pulses, this dark-induced OFF response has a similar amplitude, but a significantly shorter latency (Fig. 3d). Similar recordings indicate that cha-lOLP does not respond to dark pulses and only generates the fast ON response to light pulses.

Together, our recordings using voltage and calcium indicators demonstrates that the ON and OFF selectivity in the larval visual system emerges at the level of the lOLPs. We show that cha-lOLP specifically responds to light increments and therefore is ON selective, in contrast to glu-lOLP which responds to both light increments and decrements. However, given the significant temporal delay in its slow ON response, glu-lOLP displays an OFF selectivity at the temporal scale that matches the ON selectivity of cha-lOLP. This allows the pair of lOLPs to transmit light increments and decrements to downstream circuits. The functional significance of the slow ON response remains to be determined.

### The mAchR-B receptor mediates light-induced physiological responses in glu-OLP

Our study demonstrated that when stimulated with light, larval PRs depolarize cha-lOLP and hyperpolarize glu-lOLP. These physiological responses are likely mediated by differentially-expressed acetylcholine receptors (AchRs) in the lOLPs that respond to acetylcholine release from the PRs ^34, 35^. In particular, the sign-inversion, which transforms the light response in the PRs into an OFF response in glu-lOLP, is critical for generating ON and OFF selectivity in the larval visual circuit. Therefore, we sought to identify the receptor that mediates the light-induced inhibition in glu-lOLP and produces this sign-inversion.

While ionotropic nicotinic AchRs (nAchRs) are generally associated with neuronal activation, subtypes of muscarinic AchRs (mAchRs) can be either excitatory or inhibitory depending on the G protein coupled with the receptors. Studies in mammalian mAchRs indicate that the excitatory M1/3 types are coupled to Gαq/11, whereas the inhibitory M2/4 types are coupled to Gαi/o ^34^. The *Drosophila* genome contains three mAchRs, with type A and C coupling to Gαq/11 and type B coupling to Gαi/o ^36, 37^. Additionally, R72E03-Gal4, the enhancer Gal4 line labeling glu-lOLP, was generated using an upstream enhancer element identified in the *Drosophila* mAchR-B gene ^25, 26^, thus suggesting its expression in glu-lOLP.

Given that the mAchR-B is the most likely candidate for mediating light-induced inhibition in glu-lOLP, we further examined its expression pattern using a gene-trap line, in which a Gal4-DBD element was inserted into an intron within the 5’UTR region of the mAchR-B gene by the MiMIC transposon-mediated cassette exchange technique ^38, 39^ (Fig. 4a). The mAchR-B enhancer-driven EGFP expression revealed its extensive distribution in the 3^rd^ instar larval brain and in neurons with soma positions and projection patterns that resemble the OLPs. Immunohistochemical studies using anti-ChAT and anti-VGluT antibodies confirmed that the mAchR-B receptor expresses in glu-lOLP but not in cha-lOLP (Fig. 4b, c).

**Figure 4.**
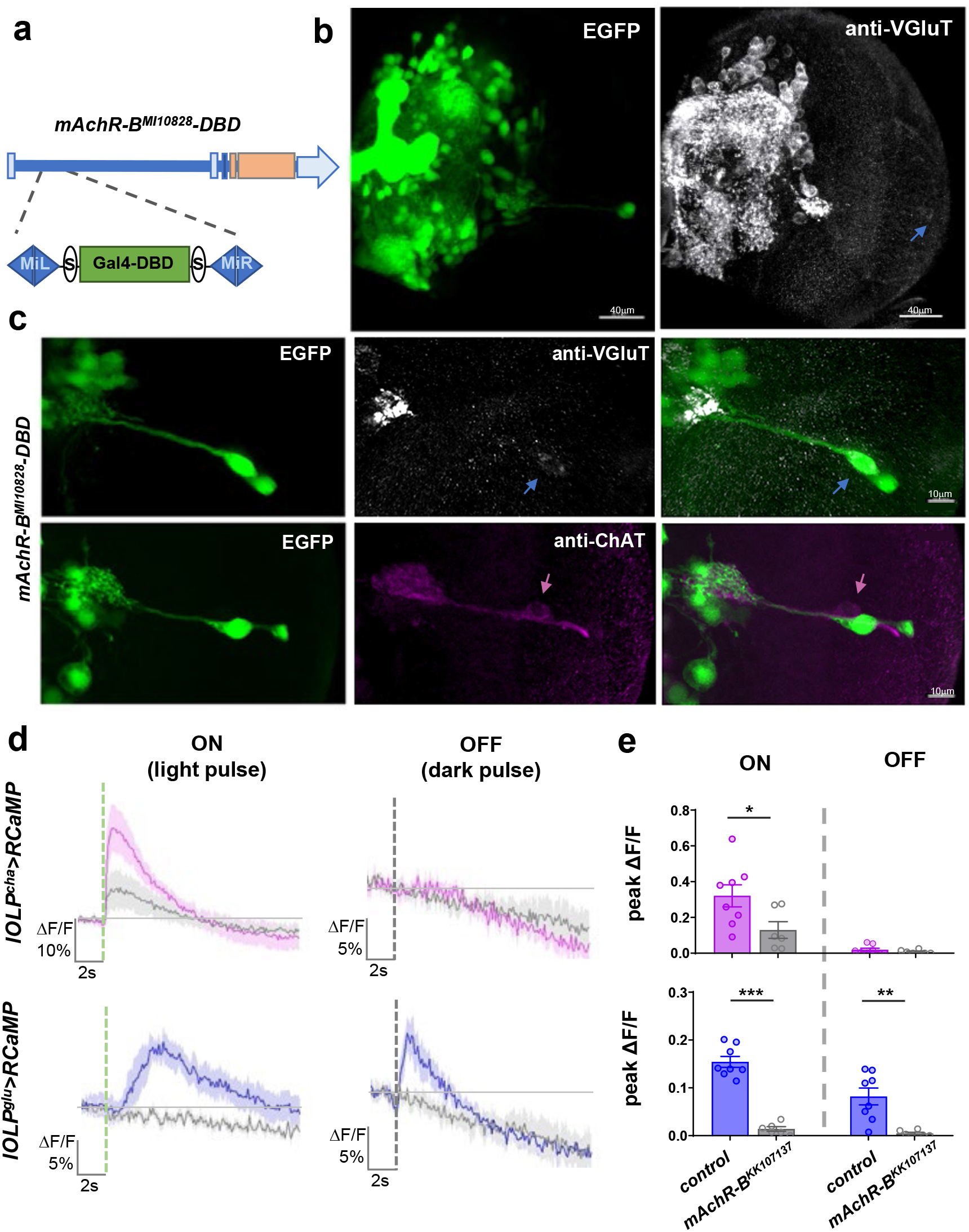
Light-induced inhibition of glu-OLP is mediated by the mAchR-B receptor. (**a**) Schematic diagram illustrating the insertion of a Gal4-DBD element into the 5’UTR region of the mAchR-B gene. Orange bar: coding exons. Light blue bar: introns. (**b**) The mAchR-B enhancer line reveals broad expression of the receptor in the 3^rd^ instar larval brain. Representative projected confocal images with EGFP expression driven by the mAchR-B enhancer (green) and anti-VGluT staining (grey) are shown. Blue arrow: glu-lOLP. **(c)** mAchR-B expresses in glu-lOLP but not cha-lOLP. The mAchR-B enhancer-driven EGFP signal colocalizes with the VGluT-positive glu-lOLP (blue arrow), but not with the ChAT-positive cha-lOLP (pink arrow). Representative projected confocal images are shown. Scale bars are as indicated. **(d-e)** Expression of mAchR-B^RNAi^ dampens cha-lOLP’s ON response and eliminates both glu-lOLP’s ON and OFF responses. The dashed green and grey lines indicate the 100 ms light or dark pulse, respectively. The genotypes are as indicated. The **(d)** average traces of the changes in lOLP>RCaMP signals and **(e)** quantification of peak values of changed intensity (ΔF/F) are shown. Shaded areas on traces and error bars on quantifications represent SEM. Control, n = 8; mAchR-B^KK107137^, n = 6. cha-lOLP, ON: p = 0.0388, t = 2.320, df = 12; cha-lOLP, OFF: p = 0.3201, t = 1.037, df = 12; glu-lOLP, ON: p < 0.0001, t = 10.09, df = 12; glu-lOLP, OFF: p = 0.0028, t = 3.736, df = 12. Statistical significance determined by student’s t-test. *P* ≥ 0.05 was considered not significant, \*\*\**P* < 0.001. \*\**P* < 0.01. \**P* < 0.05.

Next, to examine mAchR-B’s function in mediating glu-lOLP’s physiological responses, we performed knock-down experiments using a transgenic RNAi line targeting mAchR-B and recorded the lOLPs’ response to 100 ms light vs. dark pulses using RCaMP. Consistent with our earlier observations, in wild-type controls, cha-lOLP only responded to the light pulse and generated a fast calcium transient, whereas glu-lOLP responded to both light and dark pulses with either a slow or a fast calcium transient. Strikingly, mAchR-B knockdown eliminated both the light and dark pulse-induced calcium transients in glu-lOLP, indicating mAchR-B as the receptor mediating the light-induced inhibition of glu-lOLP (Fig. 4d, e). Intriguingly, the mAchR-B knock-down also significantly dampened the light responses in cha-lOLP (Fig. 4d, e), suggesting that eliminating the inhibition of glu-lOLP could potentially impact cha-lOLP’s light response.

To confirm these observations, we performed additional experiments to examine light-induced calcium transients using two-photon recordings of GCaMP6f driven by lOLP^glu^-Gal4, which showed a significantly reduced response in glu-lOLP with mAchR-B knock-down (Fig. S6), supporting the critical role of the receptor in mediating light-induced inhibition on glu-lOLP. Together, this set of experiments indicate that both ON and OFF responses of glu-lOLP are induced by mAchR-B mediated inhibition, although the downstream effector involved in producing the calcium transients could be different.

### Blocking Gαo signaling alters the temporal profile of glu-lOLP activation

To identify the downstream G protein subunits that mediate mAchR-B signaling in glu-lOLP, we examined an RNAi line targeting Gαo, the G protein subunit coupled to mAchR-B. Knocking down Gαo completely eliminated the dark pulse induced OFF response (Fig. S5b, c). Unexpectedly, knocking-down Gαo also generated a distinct phenotype in the light-induced calcium response in glu-lOLP; in addition to reducing the amplitude, Gαo knock-down also eliminated the initial reduction of calcium signal, producing a fast calcium transient instead of the typical slow ON response (Fig. 5a, b). To further characterize these effects, we genetically blocked Gαo activity using Pertussis toxin (PTX), which is previously shown to specifically inhibit Gαo in *Drosophila* ^40^. We found that expressing a transgene of PTX in glu-lOLP led to a consistent reduction in the latency of light-induced calcium responses without significantly affecting the amplitude, an effect observable in both soma and terminal regions of glu-lOLP (Fig. 5c, d).

**Figure 5.**
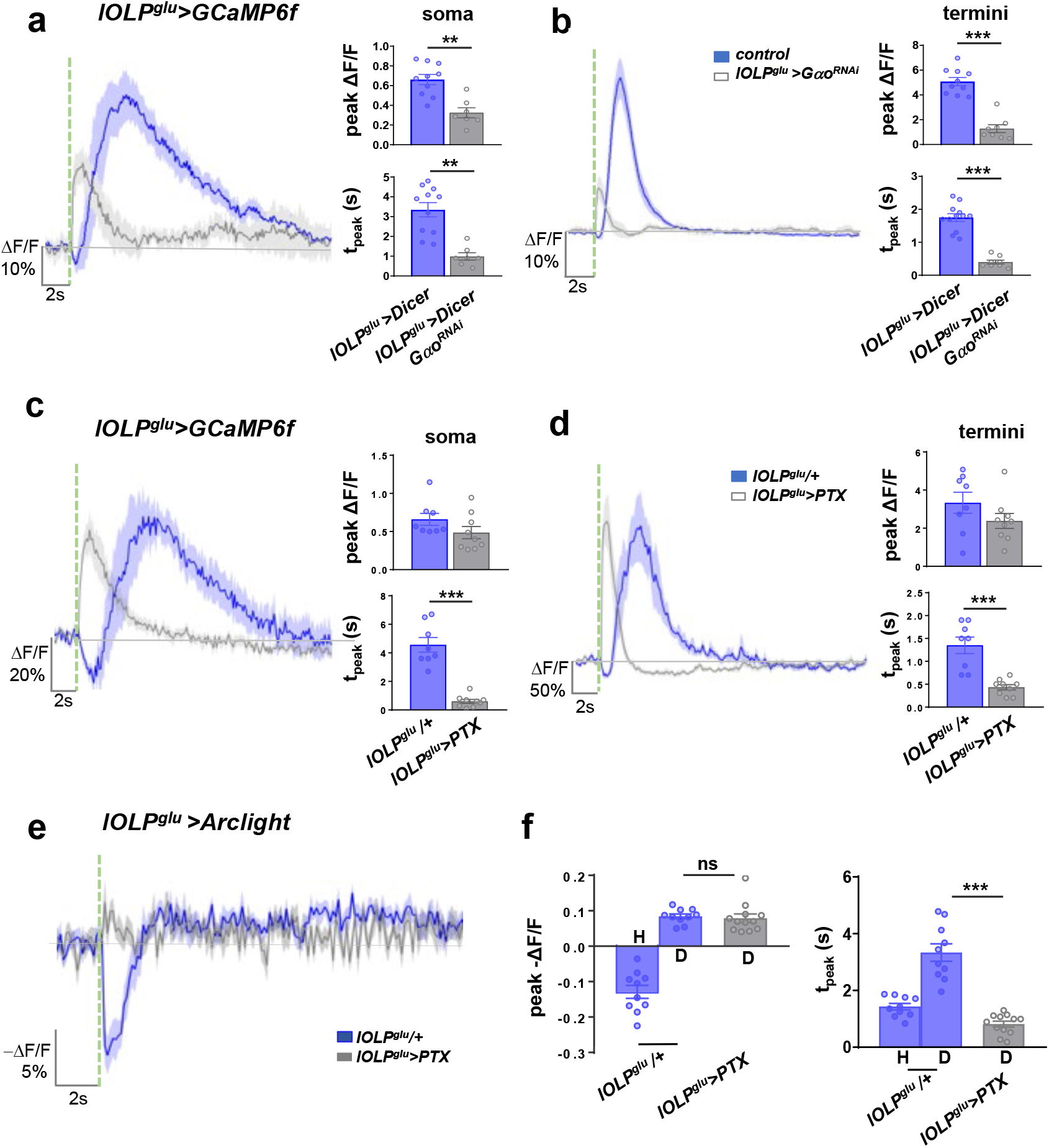
Gαo signaling regulates the temporal profile of light-evoked responses in glu-lOLP. **(a-b)** RNAi knockdown of Gαo reduced the amplitude and latency of calcium responses in glu-lOLP in both soma and termini. Average traces of the changes in GCaMP signals (left) and the quantifications of peak value and peak time (right) of changed intensity (ΔF/F) are shown. lOLP^glu^>Dicer, n = 10 for both groups; lOLP^glu^>Dicer, Gαo^RNAi^, soma: n = 7; termini: n = 8. Soma: peak value: p = 0.0005; peak time: p = 0.0002. Termini: peak value: p < 0.0001; peak time p < 0.0001. Statistical significance determined by student’s t-test. **(c-d)** Similar to the Gαo knockdown by RNAi, expression of the Gαo inhibitor PTX eliminates the delay in light-induced calcium responses in glu-lOLP without affecting its amplitude. lOLP^glu^>GCaMP6f signals were collected at the soma and termini of glu-lOLPs. Average traces of the changes in GCaMP signals (left) and the quantifications of the peak value and peak time (right) of changed intensity (ΔF/F) are shown. lOLP^glu^/+, n = 8; lOLP^glu^>PTX, n = 9 for both groups. Soma: peak value: p = 0.145; peak time: p < 0.0001. Termini: peak value: p = 0.1723; peak time: p = 0.0001. Statistical significance determined by student’s t-test. **(e-f)** PTX expression transforms the light-induced hyperpolarization into depolarization in glu-lOLP. Light-evoked voltage changes in glu-lOLP were measured by Arclight expression driven by lOLP^glu^-Gal4, which exhibits a biphasic response, a large hyperpolarization (H) followed by a small depolarization (D), in the control group. PTX expression eliminates the hyperpolarization and reveals a depolarization, corresponding to the accelerated calcium transient. Average traces of changes in Arclight signals (left) and the quantifications of the peak value and peak time (right) of changed intensity (-ΔF/F) are shown. lOLP^glu^/+, n = 10, lOLP^glu^>PTX, n = 12. Peak value: ANOVA: p < 0.0001, F = 66.92, df = 35; lOLP^glu^/+/lOLP^glu^>PTX: p = 0.9883. Peak time: ANOVA: p < 0.001, F = 42.32, df = 35; lOLP^glu^/+/lOLP^glu^>PTX: p < 0.0001. Shaded areas on traces and error bars on quantifications represent SEM. The dashed green line represents a 100 ms light pulse at 561 nm. Statistical significance determined by oneway ANOVA with post hoc Tukey’s multiple comparison’s test. ns: *P* ≥ 0.05 was considered not significant, \*\**P* < 0.01, \*\*\**P* < 0.001.

This fast, light-induced calcium transient revealed by blocking or knocking down Gαo led us to consider the possibility that, besides mAchR-B/Gαo mediated inhibition, light induces additional physiological events that could lead to calcium increases in glu-lOLP. These events are generally masked by the initial inhibition and only become visible when mAchR-B/Gαo signaling is strongly affected. It is likely that the mAchR-B RNAi line (mAchR-B^KK107137^) we used in early experiments was not effective in knocking down the receptor and therefore did not produce the same effect. To resolve the discrepancy between the mAchR-B and Gαo knock-down phenotypes, we examined a different RNAi line targeting mAchR-B (mAchR-B^HMS05691^) and observed light-induced fast calcium transient with a significantly reduced amplitude, similar to the ones shown in Gαo knock-down experiments (Fig. S6b). By comparing the outcomes generated by different manipulations that potentially block the mAchR-B/Gαo signaling to varying degrees (Fig. 5a-d, Fig. S6), we conclude that the extent and timing of glu-lOLP’s light response are regulated by the inhibition imposed by mAchR-B/Gαo signaling.

Additionally, we performed voltage recordings using Arclight and revealed a dramatic change in glu-lOLP’s voltage responses caused by PTX expression. In the control group, we observed a biphasic voltage response in glu-lOLP induced by light stimulation, which produced a large hyperpolarization event followed by a small depolarization (Fig. 5e, f). This response is temporally correlated with the biphasic calcium transients that we observed previously which showed a small reduction in calcium concentration followed by a slow rise (Fig. 5c, d). Strikingly, the expression of PTX in glu-lOLP switched the light-induced hyperpolarization to a depolarization; this result is consistent with its role in eliminating the initial reduction of the calcium concentration and producing a fast calcium rise in glu-lOLP (Fig. 5e, f).

In addition, upon examining the voltage and calcium responses in the controls and PTX expression glu-lOLPs, we found that the intracellular calcium in glu-lOLP appears to show rectification that favors depolarization events and generates large increases in calcium concentration in contrast to hyperpolarization events that produce small reductions in the calcium signals. The half-wave rectification of intracellular calcium is also observed in adult visual interneurons in the ON and OFF pathways, where the voltage to calcium transformation generates sign-inversion or preservation, which, together with the rectification, lead to the ON selectivity in Tm3 and Mi1 and OFF selectivity in Tm1 and Tm2 ^13, 29^.

In summary, our genetic studies confirm the role of mAchR-B/Gαo signaling in mediating light-induced inhibition of glu-lOLP and reveal the complexity of glu-lOLP’s light responses, which appear to contain multiple signaling events that cooperatively regulate the direction and timing of the neuron’s physiological output. Although additional studies are needed to fully characterize these responses, we found that PTX expression in glu-lOLP eliminates its OFF response while modifying the temporal profile of its ON response and thus effectively transforms glu-lOLP into an ON selective cell. Instead of transmitting light decrements, glu-lOLP expressing PTX transmits light increments to downstream VPNs and potentially disrupts the separation of the ON and OFF channels. How would this manipulation impact the physiological and behavior output of the larval visual circuit? We addressed this question in the following experiments.

### glu-lOLP regulates both cha-lOLP and VPNs in the larval visual circuit

Using PTX expression to modify the temporal profile of glu-lOLP’s activation, we examined how glu-lOLP interacts with cha-lOLP and two types of projection neurons. We expressed PTX in glu-lOLP using lOLP^glu^-Gal4 and monitored the light-induced calcium responses in all three OLPs through OLP-LexA driving expression of GCaMP6f. Consistent with our earlier observations, PTX expression accelerated the light response in glu-lOLP without affecting its amplitude. Importantly, this fast activation of glu-lOLP led to significant reductions in light-induced calcium responses in cha-lOLP (Fig. 6a, b), suggesting that glu-lOLP acts as an inhibitory input to cha-lOLP and that disrupting the temporal separation between the cha- and glu-lOLP’s light responses affects the ability of cha-lOLP to respond to light. Collectively, the direct synaptic interactions between the two lOLPs demonstrated by the connectome study ^15^, the inhibitory effect of cholinergic inputs on glu-lOLP (Fig. 3a, b) and the dampened light response in cha-lOLP generated by early activation of glu-lOLP (Fig. 6a, b) support a model of reciprocal inhibitory interactions between glu-lOLP and cha-lOLP.

**Figure 6.**
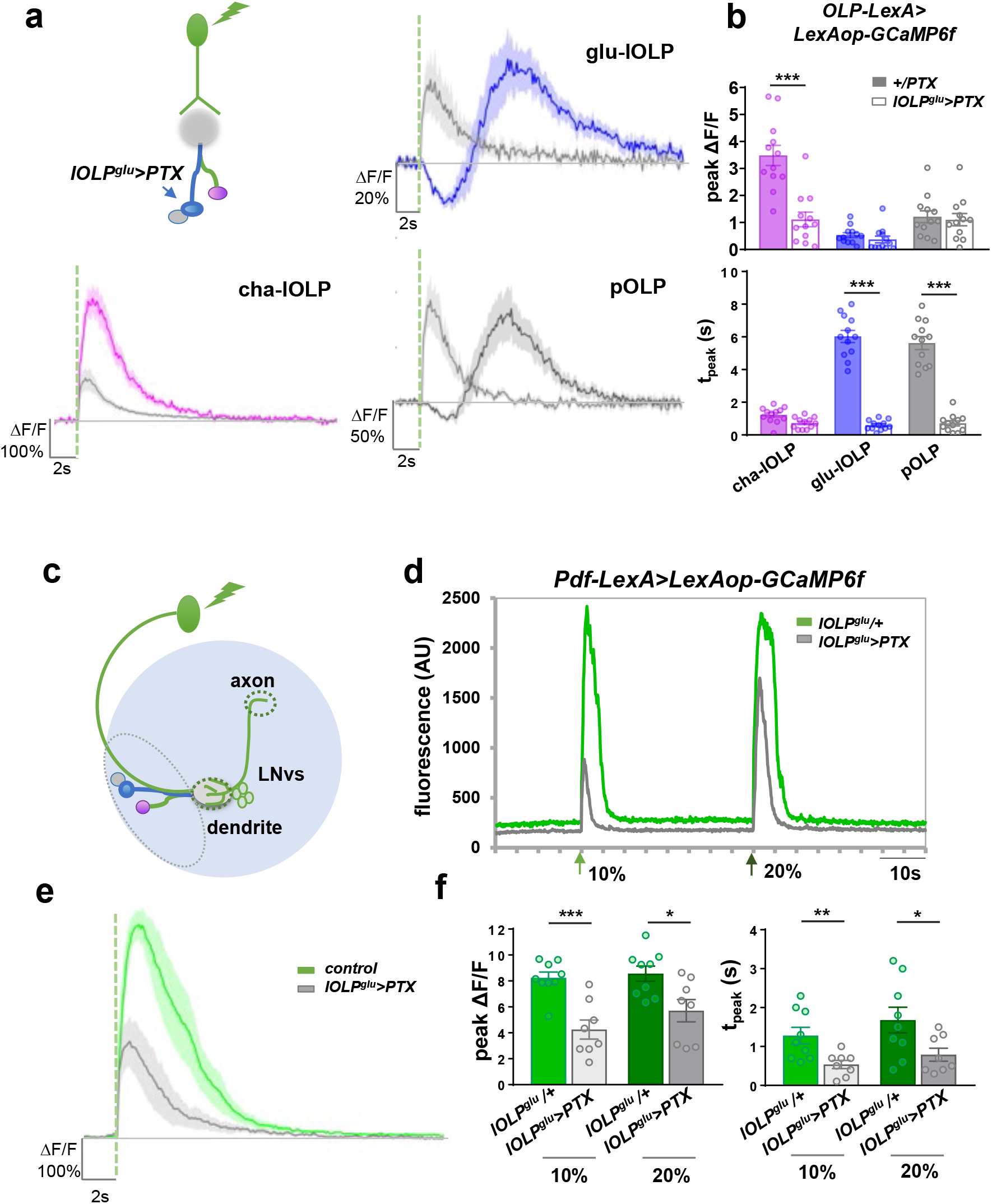
Glu-lOLP regulates light-evoked responses in cha-lOLP, pOLP and LNvs. (**a-b**) glu-lOLP inhibits cha-lOLP and activates pOLP. **(a)** Schematic diagram illustrating the experimental design, in which PTX expression is restricted to glu-lOLP while the light responses of all three OLPs are reported by OLP-LexA-driven LexAop-GCaMP6f. PTX expression eliminates the temporal delay in glu-lOLP’s light responses and leads to the dampened response in cha-lOLP and the accelerated response in pOLP. Average traces of changes in GCaMP signals are shown. The dashed green line represents a 100 ms light pulse at 561 nm. Shaded areas on traces represent SEM. **(b)** Quantifications of the peak value and peak time of changed intensity (ΔF/F) of GCaMP6f are shown. n = 12 in all groups. Peak values: cha-lOLP: p < 0.001; glu-OLP: p = 0.9967; pOLP: p = 0.9995. Peak times: cha-lOLP: p = 0.6956; glu-lOLP: p < 0.001; pOLP: p < 0.001. Error bars on quantifications represent SEM. Statistical significance determined by one-way ANOVA with post hoc Tukey’s multiple comparison’s test. (**c**) Schematic diagram showing the optical recording of light-induced responses in LNvs. Green light (561 nm) activates Rh6-PRs and elicits calcium transients in the LNvs. Pdf-LexA driven GCaMP6f signals are recorded in the axon terminal region (dashed circle). **(d)** A representative raw trace of a light-induced calcium transient in LNvs is shown. 100 ms light stimulations (green arrows) were delivered with either 10% or 20% laser power and induced robust calcium transients in LNvs. Compared to the controls, PTX expression in glu-lOLP (lOLP^glu^>PTX) leads to dampened responses with reduced durations. **(e-f)** PTX expression in glu-lOLP reduced the light-induced calcium response in LNvs. **(e)** Average traces of changes in GCaMP6f signal are shown. The dashed green line represents a 100 ms light stimulation at 10% intensity. **(f)** Quantifications of the peak value (left) and peak time (right) of changed intensity (ΔF/F) of GCaMP6f signals in LNvs, which showed significant reductions with the PTX expression in glu-OLP. Two different intensities of light stimulation generated similar results. Control: n = 9; lOLP^glu^>PTX: n = 8. Peak values: 10%: p = 0.0003, t = 4.758, df = 15; 20%: p = 0.0136, t = 2.795, df = 15. Peak times: 10%: p = 0.0090, t = 2.998, df = 15; 20%: p = 0.0354. t = 2.311, df = 15. Statistical significance determined by student’s t-test. \**P* < 0.05, \*\**P* < 0.01, \*\*\**P* < 0.001.

Blocking Gαo signaling in glu-lOLP also revealed close interactions between pOLP and glu-lOLP. PTX expression in glu-lOLP significantly reduced the latency of the light-induced response in pOLP without affecting its amplitude (Fig. 6a, b). Because of these matching temporal profiles, both with or without the PTX expression in glu-lOLP, we concluded that light-induced calcium responses in pOLP are driven by glu-lOLP’s activities (Fig. 2a, b, 6a, b, Fig. S3b). Although the connectome study did not find direct synaptic interactions between the pair, it is possible that this effect is indirect. However, the close physical proximity between glu-lOLP and pOLP also suggests that they may interact through gap junctions ^15^.

Next, we examined the impact of altered glu-lOLP kinetics on an additional group of VPNs, the larval ventral lateral neurons (PDF-LaNvs or LNvs). LNvs are a group of four peptidergic neurons that regulate the circadian rhythm in both larval and adult *Drosophila* ^41, 42^. Besides receiving synaptic inputs from the lOLPs, LNvs are also contacted directly by both Rh5- and Rh6-PRs in the LON (Fig. 6c, Fig. S7) ^15^ and are activated by cholinergic inputs mediated through nAchR signaling ^43^. Additionally, previous studies demonstrated that glutamatergic input inhibits larval LNvs through the action of a glutamate-gated chloride channel, GluCl^- 44^.

Using an LNv-specific enhancer Pdf-LexA, we expressed GCaMP6f in LNvs and recorded robust light-elicited calcium responses in the LNvs’ axon terminal region ^27^ (Fig. 6d). Importantly, expressing PTX in glu-lOLP significantly reduced both the amplitude and the duration of these calcium transients (Fig. 6d-f), suggesting glu-lOLP also provides inhibitory inputs onto the LNvs and that changing the temporal profile of glu-lOLP influences LNvs’ physiological responses to light.

Together, our results show that altering the temporal kinetics of a single neuron, glu-lOLP, strongly influences light responses in both visual interneurons and projection neurons, supporting the functional significance of the temporal control of glutamatergic transmission in the larval visual circuit. In addition, our studies also validated reciprocal interactions between cha- and glu-lOLPs and demonstrated the ability of glu-lOLP to elicit distinct physiological responses in different types of VPNs.

### glu-lOLP is required for dark-induced behavioral responses

To illustrate the potential roles for lOLPs in transmitting ON and OFF signals from the PRs to the VPNs and based on our findings and the connectivity map, we propose a model with three components. First, the pair of lOLPs exhibit distinct responses to light increments and decrements, thus acting as ON and OFF detectors. The sign-inversion required for OFF detection in glu-OLP is mediated by the mAchR-B receptor. Second, while cha-lOLP displays clear ON selectivity, glu-lOLP has both ON and OFF responses. The OFF selectivity in glu-lOLP emerges from the temporal control of its activity by mAchR-B/Gαo signaling. Third, extending our findings in the LNvs and pOLP to the rest of the VPNs, we propose that, although downstream VPNs receive both cholinergic and glutamatergic inputs, there are specific groups of ON vs. OFF-responsive VPNs that are functionally separated based on their molecular compositions. ON-responsive VPNs (ON-VPNs), such as LNvs, are activated by cholinergic signaling and inhibited by glutamatergic signaling, while OFF-responsive VPNs (OFF-VPNs), such as pOLP, behave the opposite way. Although additional physiological studies on other VPNs are needed to validate this model, given the lack of anatomical segregation of ON and OFF pathways, this functional separation of VPNs is a plausible solution to preserve and transmit the ON and OFF signals at the level of VPNs (Fig. 7a).

**Figure 7.**
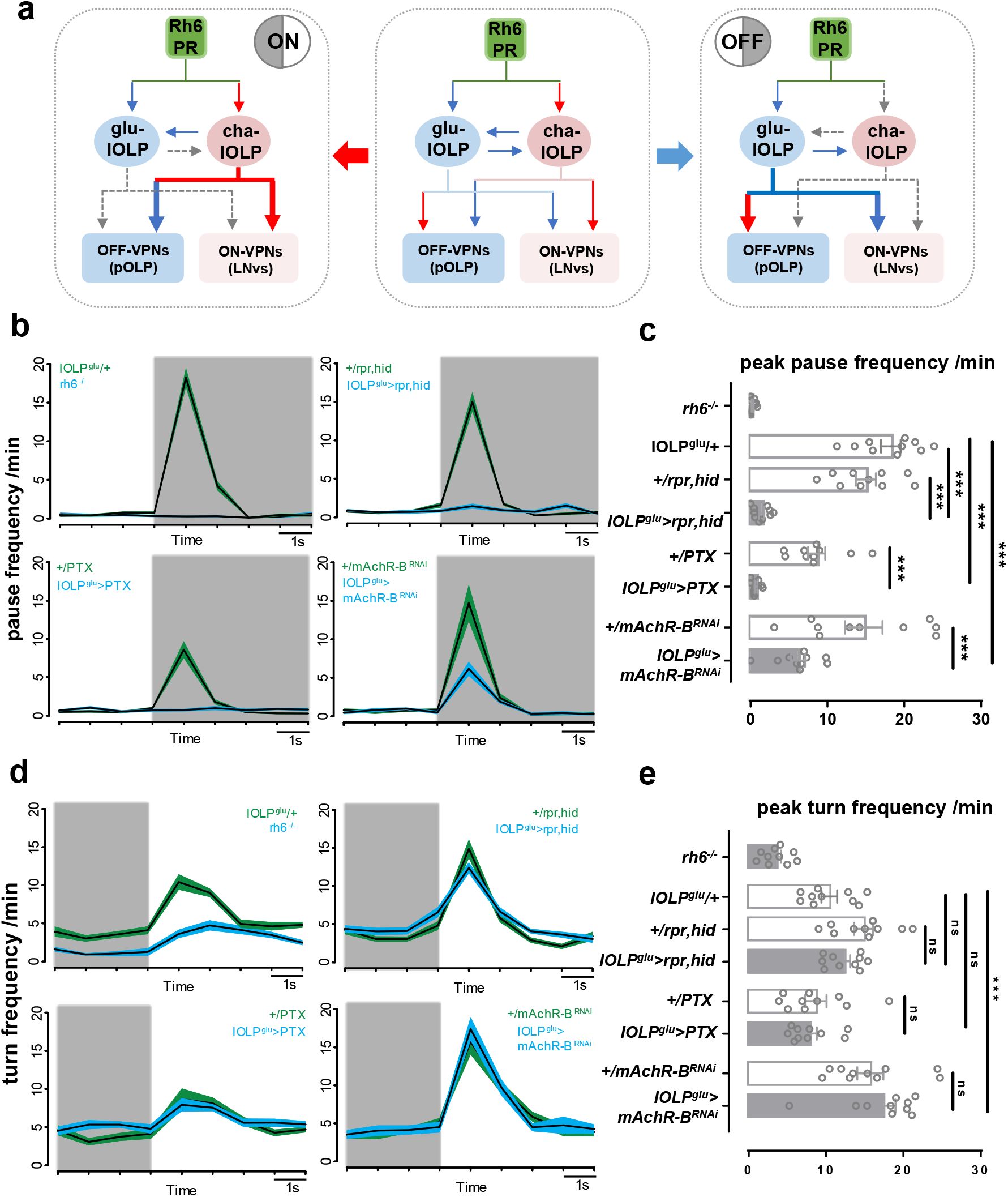
Glu-lOLP is required for dark-induced pausing behavior. **(a)** A proposed model illustrating the emergence of ON and OFF selectivity in the larval visual circuits. Middle: lOLPs detect and transmit the ON and OFF signals in the larval visual circuit. Light induces ACh release from Rh6-PRs, which activates the cha-OLP and inhibits glu-lOLP through differentially expressed AChRs. During an ON response (left panel), the cholinergic transmission is dominant, activating ON-VPNs and suppressing OFF-VPNs. During an OFF response (right panel), glu-lOLP activates OFF-VPNs and suppresses ON-VPNs. Predicated by this model, glu-lOLP is essential in initiating the OFF response. **(b-c)** Genetic manipulations of glu-lOLP affect dark-induced pausing behavior in larvae. **(b)** Plots of average pause frequency are shown. The transition from light to dark is indicated by the shade of the area. **(c)** Quantification of dark-induced pause frequency reveals the critical role of Rh6-PRs and glu-lOLP in this behavioral response. Statistical significance determined by one-way ANOVA: p < 2e-16, F = 35.6, df = 7, 72 followed by post hoc Dunnetts’s multiple comparison’s test: lOLP^glu^/+ - lOLP^glu^> rpr, hid: p < 1e-04, t = 9.907; +/rpr,hid - lOLP^glu^> rpr, hid: p < 1e-04, t = 7.990; lOLP^glu^/+ - lOLP^glu^> PTX: p < 1e-04, t = 10.337; +/PTX - lOLP^glu^> PTX: p = 0.000108, t = 4.648; lOLP^glu^/+ - lOLP^glu^> mAChR-BRNAi: p < 1e-04, t = 7.120; +/mAChR-BRNAi - lOLP^glu^> mAChR-BRNAi: p < 1e-04, t = 5.044. \*\*\**P* < 0.001. n = 10 for each genotype. **(d-e)** The light-induced increase in turning frequency is reduced in Rh6 mutants but unaffected by glu-lOLP ablation, PTX expression or mAchR-B knock-down in glu-lOLP. **(d)** Plots of average turn frequency are shown. The transition from dark to light is indicated by the shade of the area. **(e)** Quantifications of the light-induced turn frequency reveals that glu-lOLP does not influence the behavioral response induced by the dark to light transition. Statistical significance determined by one-way ANOVA: p < 7.16e-12, F = 14.93, df = 7, 72 followed by post hoc Dunnetts’s multiple comparison’s test: lOLP^glu^/+ - lOLP^glu^> rpr, hid: p = 0.753, t = −1.159; +/rpr,hid - lOLP^glu^> rpr, hid: p = 0.538, t = 1.472; lOLP^glu^/+ - lOLP^glu^> PTX: p = 0.530, t = 1.483; +/PTX - lOLP^glu^> PTX: p = 0.996, t = 0.445; lOLP^glu^/+ - lOLP^glu^> mAChR-BRNAi: p < 0.001, t = −4.134; +/mAChR-BRNAi - lOLP^glu^> mAChR-BRNAi: p = 0.849, t = −0.996. ns *P* > 0.05, \*\*\**P* < 0.001. n = 10 for each genotype.

Predicated by this model, an ON response is dominated by cholinergic transmissions from cholinergic PRs and cha-lOLP, while the extended inhibition of glu-lOLP via mAchR-B/Gαo signaling ensures the activation of only the ON-VPNs. During an OFF response, with no cholinergic input, the glu-lOLP is solely responsible for activating OFF-VPNs. In terms of their roles in regulating behavioral output, cha-lOLP likely functions in modulating the strength and duration of the light-induced response, and glu-lOLP would be essential for initiating the dark-induced behavioral response (Fig. 7a).

To test these predications and identify the functional role of glu-lOLP using the genetic tools we established, we performed behavioral experiments to quantitatively analyze responses towards dark-light or light-dark transitions during larval navigation. Previous studies indicated that, upon encountering a reduction in light intensity at a light-dark boundary, larvae increase their pausing frequencies. On the other hand, upon sensing an increase in light intensity at a dark-light boundary, larvae increase their turning frequencies. These differential responses towards light and dark are a part of the navigation strategies larvae use for efficient negative phototaxis ^18^. Notably, these behavioral changes take place in a 2-3 s time frame, matching the temporal kinetics of lOLPs’ physiological responses towards light or dark pulses.

We found that the Rh6-PR/lOLP pathway plays a significant role in mediating these behavioral responses. Behavior tests with the Rh6 mutant showed that phototransduction mediated through Rh6-PRs is necessary for dark-induced pausing. In addition, genetic manipulations of glu-lOLP, including the expression of the cell death genes *rpr* and *hid*, the Gαo inhibitor PTX and the RNAi transgene targeting the mAchR-B receptor all generated significant reductions of dark-induced pausing behavior, whereas corresponding Gal4 and UAS control larvae showed robust dark-induced pausing (Fig. 7b, c). These results indicate that either the ablation of glu-lOLP or the blocking of mAchR-B/Gαo signaling affects the dark-induced behavioral response, supporting the critical functions of glu-lOLP and mAchR-B/Gαo signaling in mediating OFF detection.

In contrast, although Rh6 mutants also exhibit deficits in light-induced increases in turning frequency, this behavioral response to light was largely unaffected by genetic manipulations of glu-lOLP. This result demonstrates that glu-lOLP is not involved in regulating larval behavioral responses towards a dark-light transition and altering glu-lOLP’s activation does not change the basic light responsiveness of the visual circuit (Fig. 7d, e), thus validating our model predicting specific involvement of glu-lOLP in dark-induced behaviors.

Although further experiments are needed to address the behavioral relevance of cha- and glu-lOLP in regulating other visually-guided behaviors, our studies measuring dark-induced pausing behavior indicate that glu-lOLP has an essential role in mediating OFF detection in the larval visual circuit, consistent with the model developed based on the anatomical and physiological studies.

## DISCUSSION

The *Drosophila* larval visual circuit, with its small number of components and complete wiring diagram, provides a powerful model to study how specific synaptic interactions support visual computation. Built on knowledge obtained from connectome and behavioral analyses, our physiological and genetic studies revealed unique computational strategies utilized by this simple circuit for processing complex outputs. Specifically, we demonstrated the essential role of glu-lOLP, a single glutamatergic interneuron, in meditating OFF detection at both the cellular and behavior levels, identified mAchR-B/Gαo signaling as the molecular machinery regulating its physiological properties and provided evidence supporting the function of temporally controlled inhibition in generating ON vs. OFF discrimination.

### Physiological studies of larval visual interneurons using optical approaches

To analyze the physiological properties of the larval visual interneurons and projection neurons, we performed optical recordings using targeted expression of calcium and voltage indicators in larval brain explants. The imaging studies in combination with the ssTEM connectome data provide us valuable information regarding the physiological properties of and synaptic interactions among lOLPs, pOLP and LNvs. However, because the larval visual circuit has not yet been examined by electrophysiological methods, we relied on optical recordings to gain information on their physiological responses. There are clear technical limitations that we needed to take into consideration when analyzing our data, including and not limited to the properties of the voltage and calcium sensors as well as the imaging and visual stimulation protocols. Nonetheless, extensive physiological characterizations of mammalian retinae and adult *Drosophila* visual ganglia provided critical references that facilitated our interpretation ^13, 29^.

To process light and dark information in parallel, both mammalian and adult fly visual systems utilize anatomical segregation to reinforce split ON and OFF pathways ^45^. In the larval visual circuit, however, almost all VPNs receive direct inputs from both cha-lOLP and glu-lOLP as well as the Rh5-PRs ^15^. Therefore, the response signs of the VPNs cannot be predicted by their anatomical connectivity to ON and OFF detectors. Furthermore, because these synaptic interactions take place in a single small neuropil, it is unlikely that the ON and OFF signals are physically separated.

Based on the cumulative evidence obtained through genetic, anatomical and physiological studies, we propose that temporal control of inhibition could potentially solve these challenges and act as the main mechanism that mediates ON vs. OFF discrimination in larvae. While cha-lOLP displays clear ON selectivity, the OFF selectivity in glu-lOLP is achieved by the extended suppression of its ON response by mAchR-B/Gαo signaling. In addition, in mammalian sensory systems, similar temporal delay of the input-evoked inhibition relative to excitation sharpens the tuning of principle cells to preferred stimuli (reviewed in ^46^). Although the temporal scale of this delay in larval lOLPs is on the order of seconds, rather than the milliseconds that are observed in mammalian cortical neurons, it is conceivable that the temporal delay in the glu-lOLP’s ON response produces a window of heightened responsiveness in cha-lOLP and ON-VPNs towards light signals. Thus, the temporal separation between cholinergic and glutamatergic transmission could reinforce the functional segregation in the VPNs and lead to distinct transmission of ON and OFF signals. Although the underlying physiological events that lead to this temporal segregation remain unclear and further validation of its function using physiological and behavioral approaches are needed, the temporal segregation of ON and OFF pathways in larval visual circuit provides an elegant solution that may be of general use in similar contexts where parallel processing is required without anatomically split pathways.

### Candidate ON- and OFF-VPNs and motion detection

The connectome study identified ten larval VPNs which receive both direct and filtered inputs from two types of PRs and transmit visual information to higher brain regions ^15^, including four LNvs (PDF-LaNs), 5^th^ LaN, nc-LaN1 and 2, pVL09, VPLN and pOLP. Previous behavioral studies suggested that subpopulations of VPNs might code distinct visual features. Although the physiological roles of these neurons in visual processing are still unknown ^17, 18, 47^, based on our findings, we expect to observe functional diversity in individual VPNs generated either by differential expression of neurotransmitter receptors or molecules involved in electric coupling.

In this study, we specifically analyzed light-induced responses in pOLP and LNvs and propose them as examples of OFF- and ON-VPNs, respectively. pOLP is driven by glu-lOLP, possibly through gap junctions, and exhibits the properties of an OFF-VPN by preserving OFF signals. At the same time, optical recordings in LNvs allow us to characterize the differential effect of cholinergic and glutamate transmission in the ON-VPNs. However, previous studies using several behavior paradigms failed to detect clear requirements for either LNvs or pOLP in larval visually-guided behaviors ^17, 18, 47^. Therefore, further physiological studies of the rest of the VPNs, the expression profiling of their neurotransmitter receptors and additional behavioral experiments targeting specific visual tasks are needed to gain a comprehensive understanding of the visual computation at the level of VPNs.

Additionally, besides the ON and OFF types that are required for basic ON vs. OFF discrimination, VPNs are also involved in encoding visual information that guides complex behaviors. The temporal regulation of their glutamatergic and cholinergic inputs as well as the possibility of local computation generated by synaptic interactions within the LON are among the possible ways to increase the functional diversity and power of VPNs in processing complex visual information. In the adult fly visual system and the vertebrate retina, ON and OFF pathways are directly linked to motion detection ^12, 48, 49^. Given the strong behavioral evidence supporting larvae’s ability to detect movements, genetic and physiological studies of the VPNs would provide useful molecular insights into motion processing ^19, 20^.

### Functional and molecular convergence of visual interneurons

Besides the similarities observed between larval lOLPs and the visual interneurons in the adult fly visual ganglia, we can also draw a close analogy between lOLPs and interneurons in mammalian retinae based on their roles in visual processing. cha-lOLP and glu-lOLP carry sign-conserving or sign-inverting functions and activate ON- or OFF-VPNs, respectively, performing similar functions as the bipolar cells in mammalian retinae ^50^. At the same time, lOLPs also provide inhibitory inputs to either ON or OFF-VPNs and thus exhibit the character of inhibitory amacrine cells ^51^. The dual role of lOLPs is the key feature of larval ON and OFF selectivity, which likely evolved based on the need for parallel processing using limited cellular resources. Although performed by a far more complex system, visual computation in mammalian retinae also heavily relies on use-dependent synaptic depression, which regulates the dynamic balance between excitation and inhibition in separate pathways ^3^.

Our studies also reveal signaling pathways shared among the visual systems. Remarkably, Gαo signaling is responsible for producing sign-inversion in both larval glu-lOLP and the ON bipolar cell in mammalian retinae ^52^. Although the two visual systems are constructed using different neurochemical systems and transmit signals through either mAchR-B or mGluR6, there appears to be a convergence in the way Gαo regulates the direction of information flow during visual processing at the molecular level. In ON bipolar cells, light increments trigger Gαo deactivation, the opening of TrpM1 channels and depolarization. Although specific molecular machineries leading to light-evoked voltage and calcium responses in larval glu-lOLP have yet to be determined, Gαo is known to have functional interactions with a diverse group of signaling molecules, including potassium and calcium channels that could directly link to the light-elicited physiological changes in glu-lOLP ^53^. Genetic and physiological studies in the larval visual circuit will facilitate the discovery of these target molecules and contribute to the mechanistic understanding of visual computation.

## METHODS

### Fly strains

The following lines were used: 1. GMR72A10-LexA, BDSC: 54191; 2. LexAop-mCherry, BDSC: 52272 ; 3. ChAT-Gal4, UAS-EGFP, BDSC: 6793; 4. GMR84E12-Gal4, (no longer available at BDSC); 5. GMR72E03-Gal4, BDSC: 47445; 6. UAS-mCD8::GFP, BDSC: 5136; 7. UAS-RedStinger, BDSC: 8547; 8. UAS-GCaMP6f, BDSC: 42747; 9. Lexop-GCaMP6f, BDSC: 44277; 10. *rh6^1^;* 11. UAS-ArcLight, BDSC: 51057; 12. UAS-RCaMP, BDSC: 51928; 13. mAchR-B^MI10828^-Gal4-DBD; 14. Tub-dVP16AD, UAS-EYFP; 15. UAS-Dcr-2, BDSC: 24651; 16. mAchR-B RNAi, VDRC: KK107137; 17. Gαo RNAi: HMS01129, BDSC: 34653; 18. mAchR-B RNAi, BDSC: 67775; 19. UAS-PTX; 20. Pdf-LexA. Stock #10 is a gift from Dr. Claude Desplan. The rest of the lines were from Bloomington Stock Center (BDSC) or Vienna Drosophila Resource Center (VDRC). Stock #13 was generated using the MI10828 MiMIC insertion in the first intron of the mAchR-B gene. A gene-trap cassette containing the Gal4-DBD sequence in place of the original Gal4 sequence, was inserted into MI10828 using ΦC31 technology by Rainbow Transgenic Flies (CA) ^38, 39, 54^. Stock #14 is as described ^39^.

#### Fly culture

Fly stocks are maintained using the standard cornmeal medium in humidity controlled 25°C incubators with a 12-hour light: 12-hour dark schedule. Light intensity in the incubator is around ~1000 lux. All immunohistochemistry studies and optical imaging were performed using wandering 3^rd^ instar larvae.

### Immunohistochemistry

The procedures for dissection, fixation and immunochemistry on larval brains were as described previously ^27^. Larval brains were collected from wandering 3rd instar larvae and fixed in 4% PFA/PBS at room temperature for 30 min, followed by washing in PBST (0.3% Triton-X 100 in PBS) and incubating in the primary antibody overnight at 4°C. Brains were then washed with PBST and incubated in the secondary antibody at room temperature for 1 hour before final washes in PBST and mounting on the slide with the antifade mounting solution. Primary antibodies used were rabbit anti-GFP antibody (Abcam, Ab6556, 1: 200), mouse anti-ChAT (DSHB, ChAT4B1, 1:10) and rabbit anti-VGluT (a gift from Dr. DiAntonio, 1:5000). Secondary antibodies used were goat anti-rabbit Alex 633 (Invitrogen, A-21070) and donkey anti-mouse CY3 (Jackson ImmunoResearch Labs, 715165150). Wholemount brain samples were treated and mounted on slides using the SlowFade Antifade kit (Life Technologies S2828).

### Confocal and two-photon imaging

Fixed samples were imaged on a Zeiss 700 confocal microscope with a 40x oil objective. Serial optical sections were obtained from whole-mount larval brains with a typical resolution of 1024 μm x 1024 μm x 0.5 μm. Two-photon imaging of genetically encoded sensors, including GCaMP6f and Arclight, was performed on a Zeiss LSM 780 confocal microscope equipped with a Coherent Vision II multiphoton laser. Time-lapse live imaging series were acquired at 100 ms per frame for 1000 frames using a 40x water objective with the two-photon laser tuned to 920 nm. Typical resolution for a single optical section is 256μm x 96 μm with 3x optical zoom. RCaMP signals were collected with similar optical and temporal resolutions, using either the two-photon laser tuned to 1040 nm (Fig. 3b) or the confocal laser at 561 nm (Fig. 3c, d, 4d, Fig. S5b, S8).

### Visual stimulation

All optical recordings, except for the experiments described above, were collected using the two-photon laser. The preparation was stimulated by 100 ms light pulses. The blue light (488 nm) or the green light stimulation (561 nm) is produced by an Argon multiline laser set at 488 nm or a DPSS-561 nm laser, respectively. Both lasers are incorporated into the LSM780 confocal microscope and controlled by the photobleaching program in the Zen software. The spectral sensitivity of *Drosophila* Rh5 and Rh6 have been previously established ^30^. Rh6 detects light within the 400-600 nm range and its maximal spectral sensitivity is ~437 nm, while Rh5 detects light from 350-500 nm and its maximal spectral sensitivity is ~508 nm. Therefore, Rh6-PRs are sensitive to light stimulations at both 488 nm and 561 nm, whereas Rh5-PRs only respond to blue light at 488 nm.

The intensity of the light stimulation was adjusted by the power setting of the laser. As measured by a light meter (Thorlabs, Germany, Model: PM100D) equipped with a light sensor (Thorlabs, Germany, Model: S170C), the output was approximately 39 μW/cm^2^ for the 561 nm laser and 11.7 μW/cm^2^ for the 488 nm laser at 20% laser power. At 10% laser power, the output was around 21.5 μW/cm^2^ for the 561 nm laser and 5.9 μW/cm^2^ for the 488 nm laser. During a 1000 frame recording collected at 100 ms per frame, two separate light pulses of different wavelengths (488 nm vs. 561 nm) or different intensities (10% vs. 20% laser power) were delivered at the 200^th^ and 600^th^ frames (Fig. S3a).

To study the responses of lOLPs to the onset and offset of extended light exposures (Fig. 3c), we collected RCaMP signals using the 561 nm confocal laser with the power setting of 0.5%, while tuning the light cycle using the 488 nm laser with the power setting of 5%. The laser power output during the light exposure was ~ 3.9 μW/cm^2^. When the 488 nm laser was turned off, the output was reduced to ~1 μW/cm^2^.

To measure the ON response using confocal recording of RCaMP (Fig. 3d, 4d, Fig. S6) (the response of lOLPs to light pulses), we collected RCaMP signals using the 561 nm confocal laser with the power setting of 0.5%-1%, while stimulating the preparation using a 100 ms light pulse generated by the 488 nm laser with the power setting of 20%. The laser power during the recording was ~ 1-2 μW/cm^2^ and increased to ~ 12.5 μW/cm^2^ with the light pulse.

To measure the OFF response (Fig. 3d, 4d, Fig. S5) (the response of lOLP towards light decrements), we recorded RCaMP signals using the confocal laser at 561 nm with the power setting of 5% plus additional illumination using the confocal laser at 488 nm with the power setting of 2%, which produced an output of ~ 11.7 μW/cm^2^. The 100 ms dark pulse was delivered by the photobleaching program with no laser activated and therefore produced a reduction of light intensity from ~ 11.7 μW/cm^2^ to 0.

### Larval eye-brain explant preparation for live imaging

Optical recordings were performed on explant preparations collected during the subjective day between ZT1-ZT8 (ZT: zeitgeber time in a 12:12 h light dark cycle; lights-on at ZT0, lights-off at ZT12). Procedures for dissection and preparation of larval brain explants were as described ^27^. The eye-brain explant containing the Bolwig’s organ, the Bolwig’s nerve, eye discs and the larval brain were dissected in PBS. The explant was carefully separated from the rest of the larval tissue without damaging the optic nerve or brain lobes, transferred into an external saline solution (120 mM NaCl, 4 mM MgCl_2_, 3 mM KCl, 10 mM NaHCO_3_, 10 mM Glucose, 10 mM Sucrose, 5 mM TES, 10 mM HEPES, 2 mM Ca2+, PH 7.2) and maintained in a chamber between the slide and cover-glass during the recording sessions.

### Imaging data analysis

Time lapse imaging series were first processed using the Zen software. Regions of interest (ROIs) around individual soma or the terminal processes were manually selected for each sample. Examples of raw images of optical recordings using OLP>GCaMP6f with the ROI selection are shown in Fig. S3b. A txt. file containing the intensity value of each ROI for individual frames within the time series was generated by the Zen software and exported to be further processed in MATLAB. No averaging, normalization or bleaching correction was performed on the imaging data set.

The quantification and graphing of the imaging data were performed using a custom written MATLAB script. Specifically, the average fluorescence intensity of the 20 frames prior to the stimulation was computed as F0. The change of fluorescence intensity after the stimulation was computed as (Ft-F0)/F0 (ΔF/F). For each sample, the peak amplitude, defined as the highest value of ΔF/F within the 80 frames after the stimulation, and the peak time, defined as the time point when peak ΔF/F is achieved, were computed and used for statistical analyses. Most traces in figures were generated by plotting the average ΔF/F of individual samples +/− standard error of the mean for each frame for the duration of 20 sec or 200 frames using a customized MATLAB script. Results presented in Fig. 2a, b, 3c, 6d and S3 are plotted with Microsoft Excel using the raw fluorescence intensity data.

### Behavioral experiments

Preparation and performance of behavioral experiments was during the day under red light conditions. Larvae were removed from food vials and cleaned with water. For each experiment, 30 early 3^rd^ instar larvae were collected with a fine brush and dark adapted for at least 10min before the start of the experiments. The larvae were placed in the middle of the testing plate made of a petri dish (BD Falcon BioDishXL, BD Biosciences) of size 24.5 x 24.5 cm that was filled with 2% agarose (Agarose Standard, Roth). Experiments were performed in a black box illuminated with red LEDs (623 nm, Conrad). A camera (acA2500-14gm, Basler AG, Germany) equipped with a Fujinon lens (Fujinon HF12.5HA-1B 12.5 mm/1.4, Fujifilm, Switzerland) and a red bandpass filter (BP635, Midwest Optical Systems, USA) was placed on top of the arena and recorded the larval behavior for 11 min at the rate of 13 frames/s. The first min of each experiment was not used for the analysis to allow the larvae to adapt to the testing plate.

During the recording period, an ON/OFF light cycle was delivered to the larvae on the testing arena by a light source made of blue and green LEDs (PT-120, Luminus, Billerica, MA, USA). The LED lights illuminated the testing plate from the top at a height of 45 cm. The intensity was 378 *μ*W/cm^2^, with peaks at 455 nm (11.9 *μ*W/cm^2^) and 522 nm (3.7 *μ*W/cm^2^) with half-widths of 9 and 14 nm, respectively. An Arduino running a customized script was used to switch the LEDs off for 1 min and on for 1 min, repeating 5 times per experiment. For image acquisition and larval behavior analyses, customized software developed in LabVIEW and the MAGATAnalyzer were used, respectively (Gershow, M. *et al*. 2012, Kane, E. A. *et al*. 2013). MATLAB and R Studio were used for further analysis, statistics and graphing.

### Behavioral data analyses

The definitions and thresholds of the behavioral parameters were as described ^17, 18, 21^. A run was defined as an event of forward locomotion with larval head and body aligned. A turn or a pause were defined as an event of slow or no forward locomotion. The speed threshold was determined for each larva individually. An event is marked as turn/pause in cases where larval velocity is slower than the average speed directly before and after a turn/pause. The head and body were aligned during a pause and not aligned during a turn. In other words, turns possess at least one head sweep, whereas pauses do not possess head sweeps. An event is marked as a head sweep in cases where the body bend angle was greater than 20°. A head sweep ends when the body bend angle is again lower than 10°. An accepted head sweep is followed by a run and a rejected head sweep is followed by another head sweep. We calculated the pause frequency per min per animal by determining the number of pauses during a 1 sec time window, multiplying this value by 60 and dividing it by the number of larvae present in the field of view of the camera during the respective time window. The turn frequency per min per animal was calculated in the same way.

### Statistical analysis

Statistical analyses for optical recordings were performed using GraphPad Prism. The twotailed unpaired student’s *t* test was used to compare data in two groups with equal or unequal sample numbers. For data containing multiple groups, one-way ANOVA was used with post-hoc Tukey’s multiple comparison test. Data in figures are presented as mean ± SEM. *P* ≥ 0.05 was considered not significant (ns); \**P* < 0.05, \*\**P* < 0.01, \*\*\**P* < 0.001.

For behavioral experiments, the statistic functions “aov” and “glht(multcomp)” in R Studio were used for statistical analyses. One-way ANOVA followed by Dunnett’s multiple comparison test was performed. *P* ≥ 0.05 was considered not significant (ns), *** *P* < 0.001. Exact n-values, degrees of freedom, F-values, t-values and p-values are provided in the figure legends.

## List of fly genotypes by figures

**Figure 1:**

Panel b: *R72A10-LexA, LexAop-mCherry; ChAT-Gal4, UAS-EGFP*

Panel c: *UAS-mCD8::GFP; GMR84E12-Gal4, UAS-RedStinger*

*UAS-mCD8::GFP; GMR72E03-Gal4, UAS-RedStinger*

Panel d: *UAS-GCaMP6f; GMR84E12-Gal4*

*UAS-GCaMP6f; GMR72E03-Gal4*

**Figure 2:**

*Wild-type: R72A10-LexA, LexAop-GCaMP6f;* +

*rh6^−/−^: R72A10-LexA, LexAop-GCaMP6f; rh6^1^*

**Figure 3:**

Panels a and b: *UAS-Arclight; GMR84E12-Gal4, UAS-RCaMP*

Panel c: *GMR84E12-Gal4, UAS-RCaMP*

Panel d: *GMR72E03-Gal4, UAS-RCaMP*

**Figure 4:**

Panels b and c: *Tub-VP16AD, UAS-EGFP; mAchR-B^MI10828^-Gal4-DBD*

Panel d: *UAS-Dicer2, GMR84E12-Gal4, UAS-RCaMP*

*UAS-Dicer2, GMR84E12-Gal4, UAS-RCaMP, mAchRB-RNAi^KK107137^*

**Figure 5:**

Panels a and b: *UAS-GCaMP6f; GMR72E03-Gal4*

*UAS-GCaMP6f; GMR72E03-Gal4, GαoRNAi*

Panels c and d: *UAS-GCaMP6f; GMR72E03-Gal4*

*UAS-GCaMP6f; GMR72E03-Gal4, UAS-PTX*

Panels e and f: *UAS-Arclight; GMR72E03-Gal4*

*UAS-Arclight; GMR72E03-Gal4, UAS-PTX*

**Figure 6:**

Panels a and b: *R72A10-LexA, LexAop-GCaMP6f; UAS-PTX*

*R72A10-LexA, LexAop-GCaMP6f; GMR72E03-Gal4, UAS-PTX*

Panels c-f: *Pdf-LexA, LexAop-GCaMP6f; GMR72E03-Gal4*

*Pdf-LexA, LexAop-GCaMP6f; GMR72E03-Gal4, UAS-PTX*

**Figure7:**

*rh6^1^*

*GMR72E03-Gal4 UAS-rpr, UAS-hid;;*

*UAS-rpr, UAS-hid;; GMR72E03-Gal4*

*UAS-PTX*

*GMR72E03-Gal4, UAS-PTX*

*UAS-Dicer2; mAchR-B-RNAi^KK107137^*

*UAS-Dicer2; GMR72E03-Gal4, mAchR-B-RNAi^KK107137^*

**Supplementary Figure 1:**

*UAS-mCD8::GFP; GMR72E03-Gal4*

*UAS-mCD8::GFP; GMR84E12-Gal4*

*UAS-mCD8::GFP; GMR72A10-Gal4*

**Supplementary Figure 2:**

*hs-flp; UAS>CD2>mCD8::GFP, GMR84E12-Gal4*

**Supplementary Figure 3:**

*R72A10-LexA, LexAop-GCaMP6f;*

*R72A10-LexA, LexAop-GCaMP6f; GMR72E03-Gal4, UAS-PTX*

**Supplementary Figure4:**

*UAS-GCaMP6f; GMR72E03-Gal4*

*UAS-GCaMP6f; GMR84E12-Gal4*

*GMR72A10-LexA; LexAop-GCaMP6f*

**Supplementary Figure 5:**

*UAS-Dicer2, GMR84E12-Gal4, UAS-RCaMP*

*UAS-Dicer2, GMR84E12-Gal4, UAS-RCaMP, mAchRB-RNAi^KK107137^*

*UAS-Dicer2, GMR84E12-Gal4, UAS-RCaMP, GαoRNAi^HMS01129^*

**Supplementary Figure 6:**

*UAS-GCaMP6f; GMR72E03-Gal4*

*UAS-GCaMP6f; GMR72E03-Gal4, mAchR-B-RNAi^KK107137^*

*UAS-Dicer2; GMR72E03-Gal4*

*UAS-Dicer2; GMR72E03-Gal4, mAchR-B-RNAi^HMS05691^*

**Supplementary Figure 7:**

*Pdf-Gal4, UAS-mCD8::GFP; R72A10-LexA, LexAop-mCherry;*

**Supplementary Figure 8:**

*UAS-GCaMP6f; GMR84E12-Gal4*

*GMR84E12-Gal4, UAS-RCaMP*

## SUPPLEMENTAL INFORMATION

Supplemental information includes 8 figures with legends.

## AUTHOR CONTRIBUTIONS

B. Q. and Q. Y. designed the experiment and performed data collection for optical recordings. T.H. performed the behavioral experiments and quantifications. A. K., H. K. and J. S. performed experiments and data analyses. F. D. generated the mAchR-B-Gal4-DBD transgenic fly. B. W. and S.S provided advice and supervision. B. Q., A. K., and Q. Y. wrote the manuscript.

## ACKNOWLEDGEMENT

We thank Mark Stopfer and Ralph Nelson for helpful discussions and comments on manuscripts. We thank C. Desplan, M. Rosbash and G. Roman for the mutant and transgenic Drosophila lines and A. DiAntonio for anti-VGluT antibody. This work was supported by the Intramural Research Program of NINDS, NIH. Grant number: NIH/ZIA-NS003137.

---

**Supplementary Figure 1.**
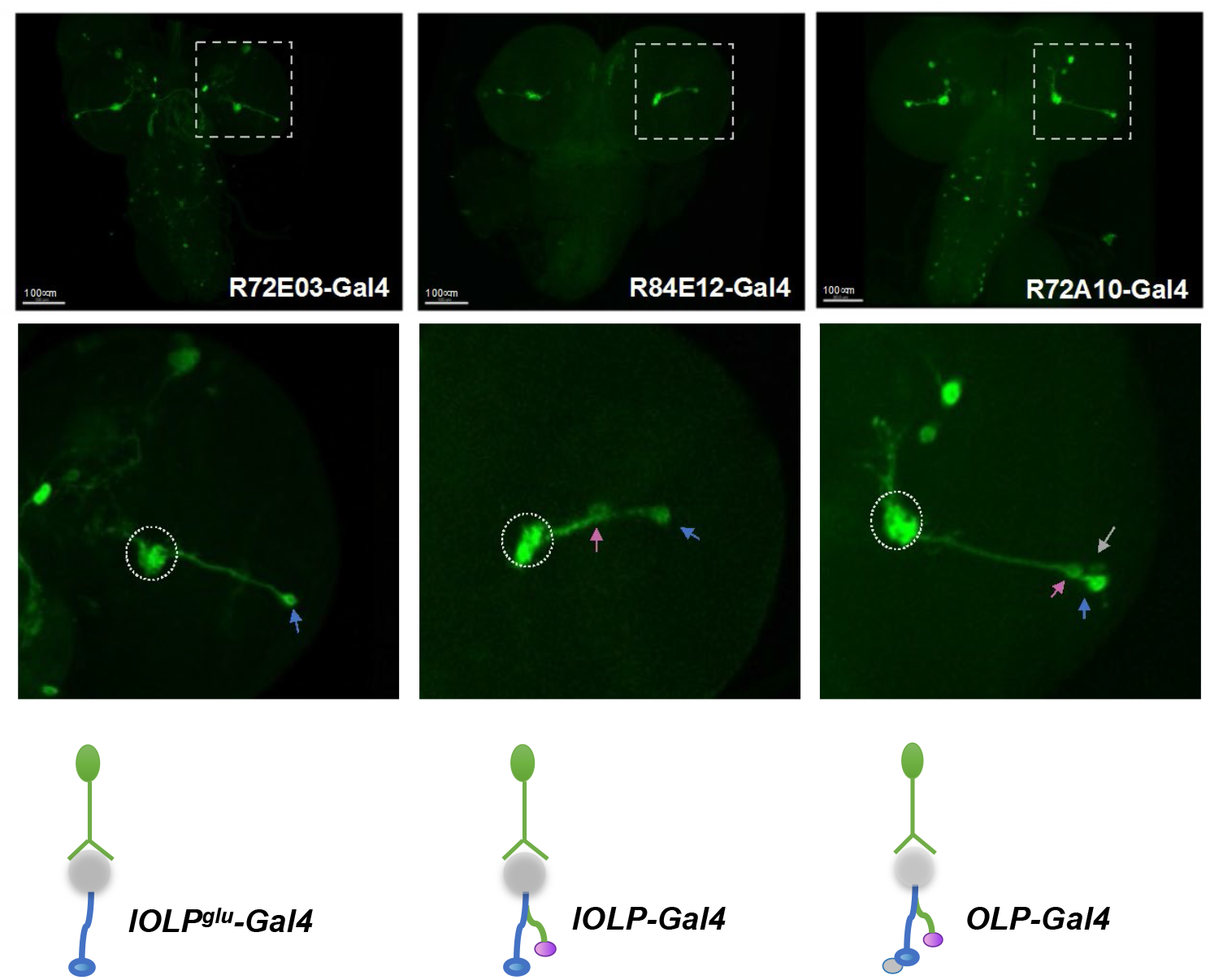
Enhancer-Gal4 lines used for labeling and manipulating the OLPs. Three Gal4 lines were selected from the Janelia Farm FlyLight Gal4 collection. Top: Representative projected confocal images showing enhancer Gal4 driven mCD8::GFP expression in larval brain tissues. Middle: zoomed-in views of the OLPs. Grey dashed circles: OLPs’ processes in the LON. Pink arrow: cha-lOLP, blue arrow: glu-lOLP, grey arrow: pOLP. Bottom: schematic illustrations showing Gal4s labeling 1, 2 and 3 OLPs. Scale bar = 100 μm. Genotypes are as indicated.

**Supplementary Figure 2.**
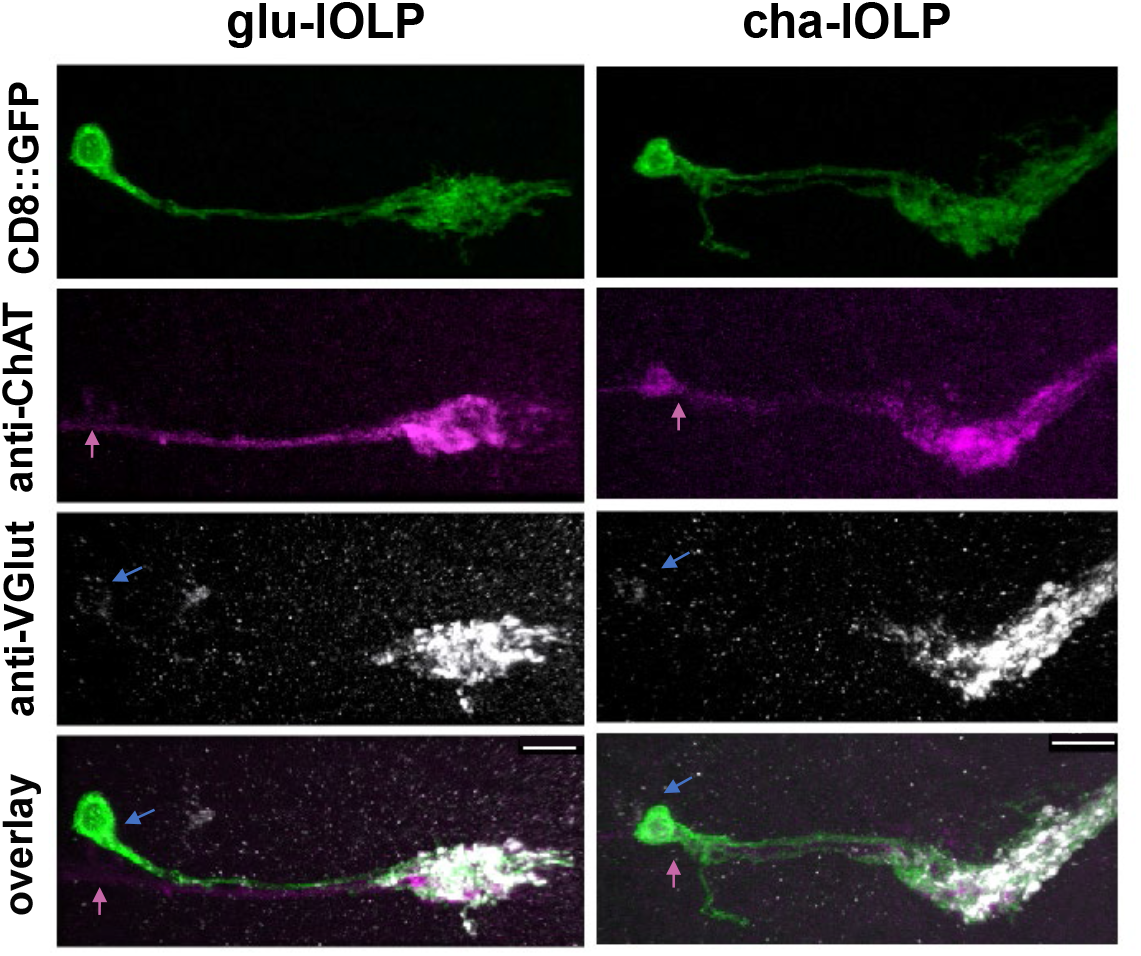
R84E12-Gal4 (lOLP-Gal4) labels glu-lOLP and cha-lOLP. Projection patterns of the two lOLPs are shown with processes terminating in the LON. The lOLPs were individually labeled by the Flip-out technique (hs-flp; lOLP-Ga4; UAS>CD2>CD8::GFP) and the identities of the lOLPs confirmed by immunostaining with anti-ChAT and anti-VGlut antibodies. Pink arrow: cha-lOLP, blue arrow: glu-lOLP. Scale bar = 20μm.

**Supplementary Figure 3.**
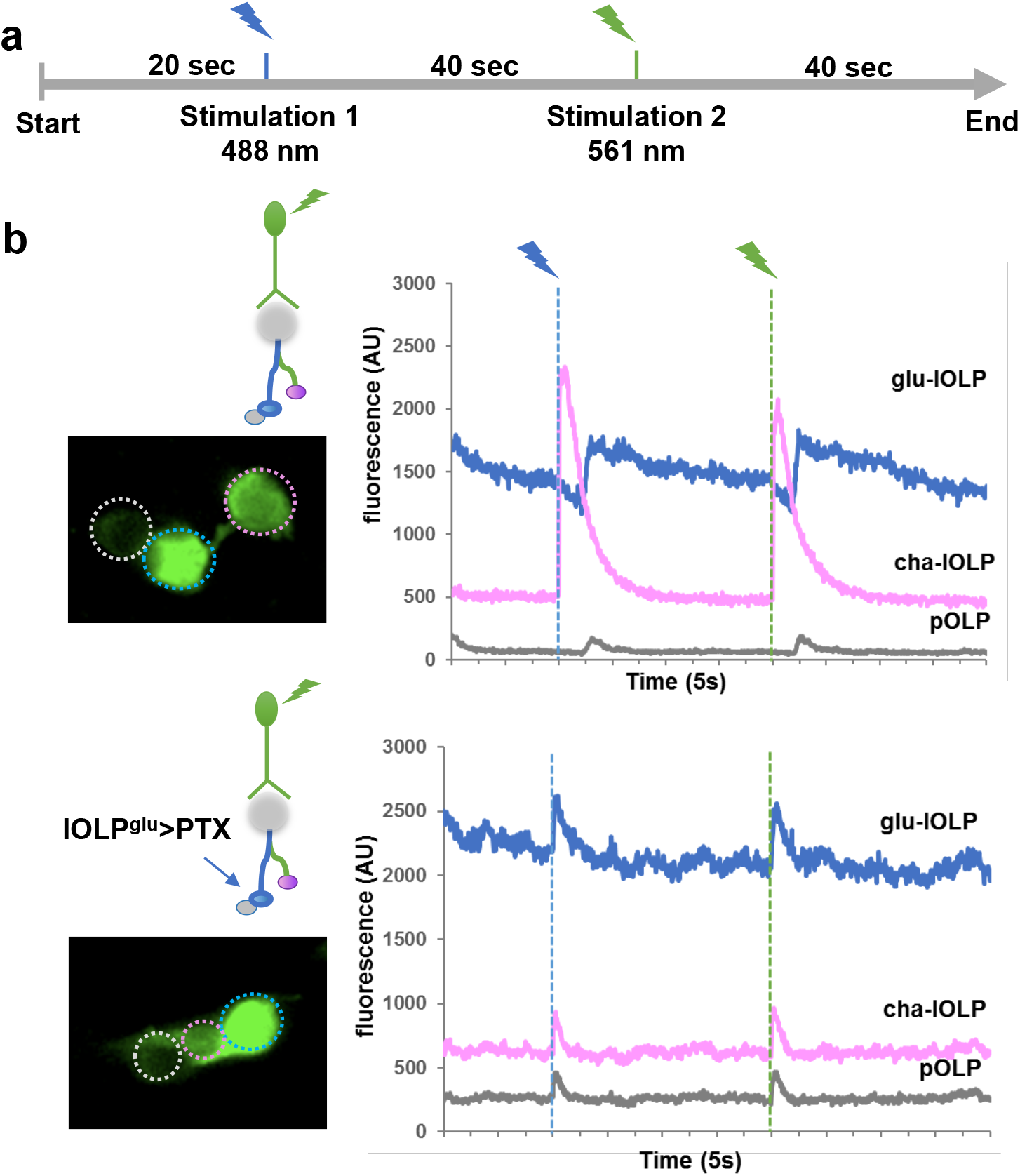
glu-lOLP inhibits cha-lOLP and activates pOLP. **(a)** Schematic illustration of a recording session with two 100 ms light pulses delivered at 488 nm (blue) or 561 nm (green). **(b)** PTX expression in glu-lOLP modifies the temporal profile of its light response and also leads to the dampened response in cha-lOLP and the accelerated response in pOLP. Representative frame and raw traces of the GCaMP recordings are shown. Top: the control with OLP-LexA driven LexAop-GCaMP6f expression. Bottom: UAS-PTX expression driven by lOLP^glu^-Gal4 was used to specifically modify the temporal profile of glu-lOLP. Somas are marked by dashed circles. Magenta: cha-lOLP, blue: glu-lOLP, grey: pOLP. Dashed lines indicate light pulses.

**Supplementary Figure 4.**
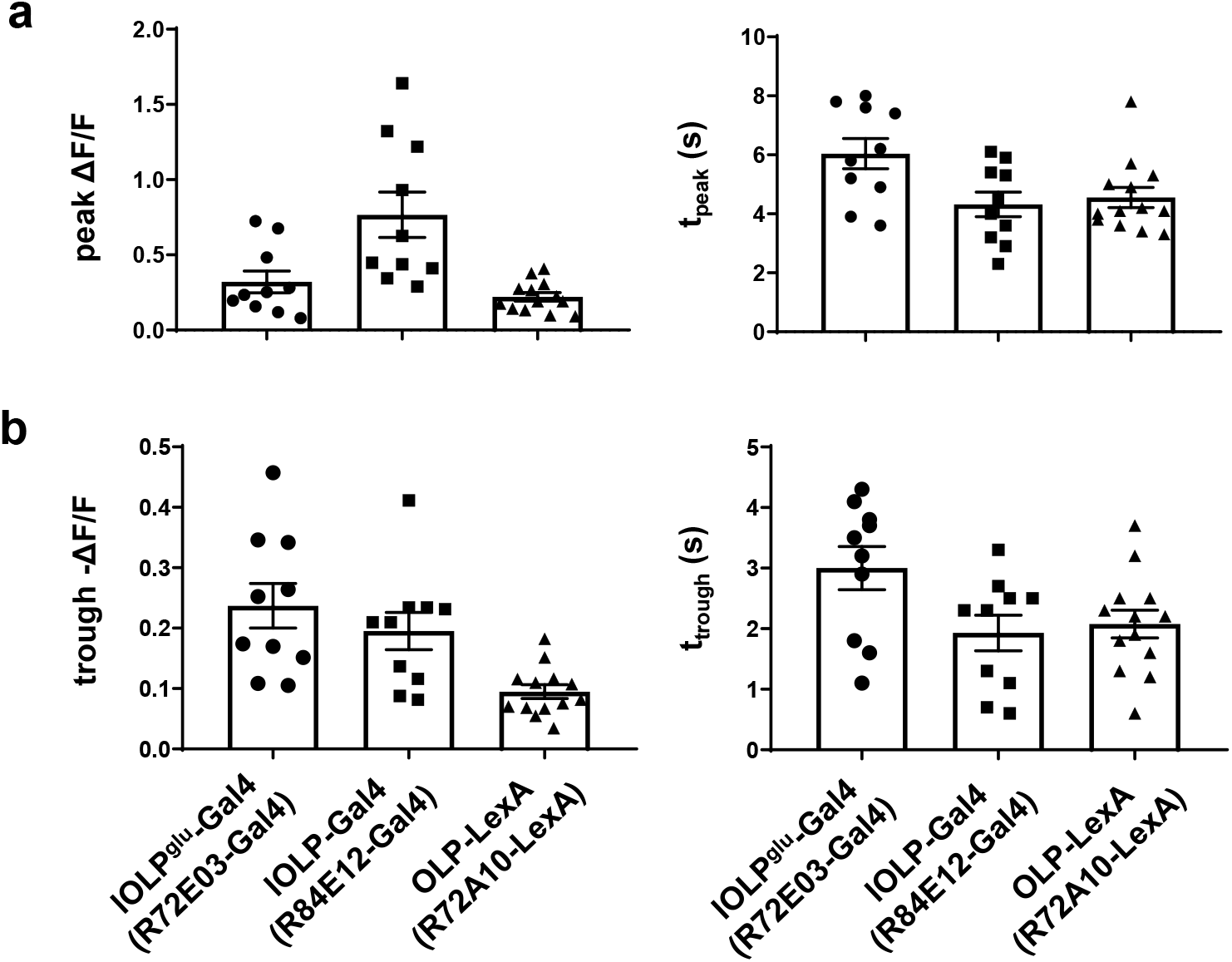
Comparison of the glu-lOLP calcium imaging results obtained using three enhancer driver lines. GCaMP6f expression in glu-lOLP driven by different enhancer lines were used to measure light induced calcium responses, which exhibited a biphasic waveform with a initial reduction followed by a slow rise of the GCaMP signals. The amplitude and latency of the peak and trough are quantified and the results obtained from three drivers are comparable. Genotypes are as indicated. Quantifications of **(a)** peak value and peak time of changed intensity (ΔF/F) are shown as well as **(b)** trough value and trough time of changed intensity (-ΔF/F). n = 10, lOLP^glu^-Gal4; n = 10, lOLP-Gal4; n = 13 OLP-LexA.

**Supplementary Figure 5.**
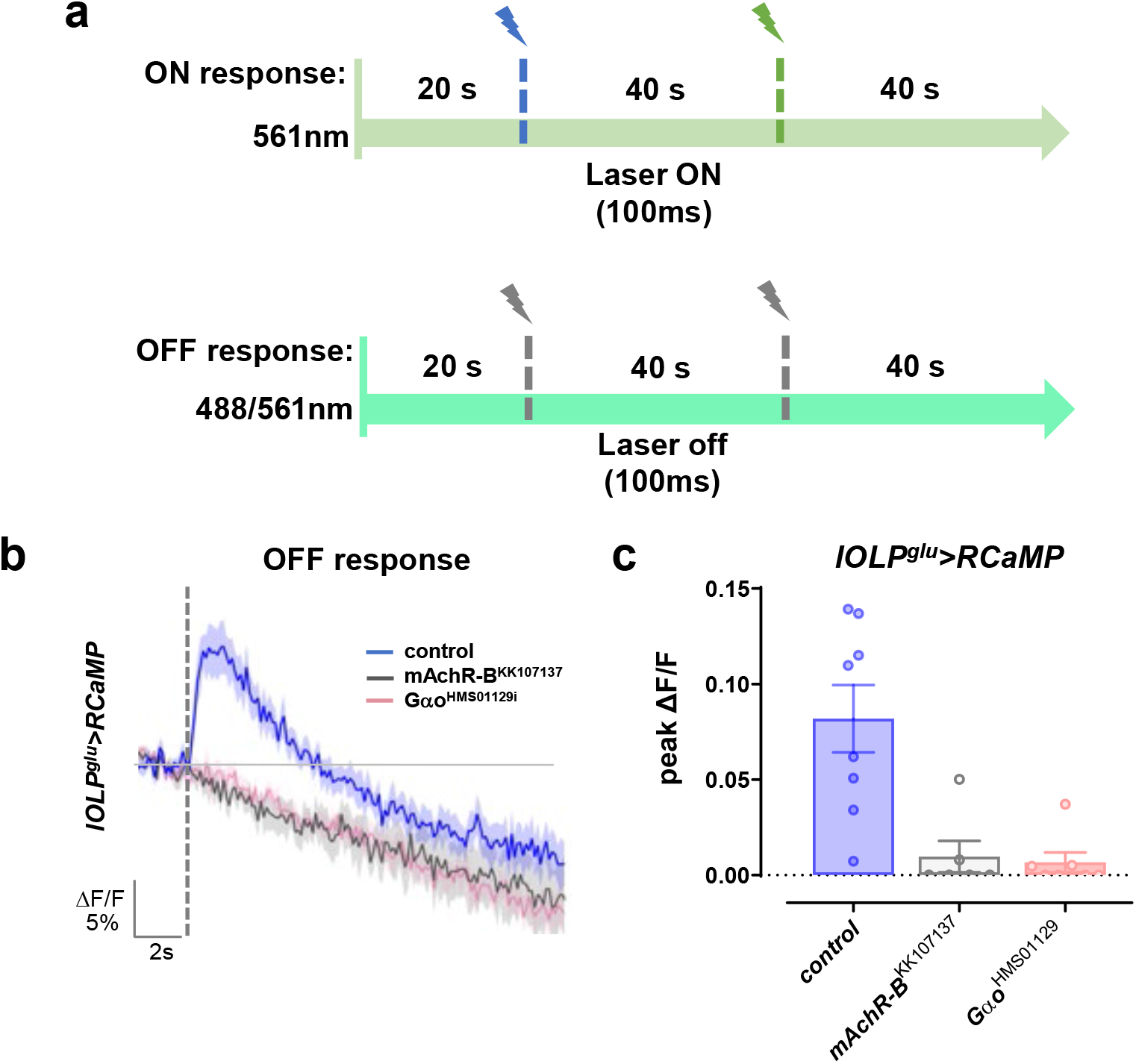
The OFF response in glu-lOLP is mediated by the mAchR-B receptor and Gαo signaling. (**a**) Schematic illustration of recordings examining the ON and OFF responses using lOLP-Gal4 driving RCaMP. The ON response was recorded using a 561 nm confocal laser at low intensity and stimulated with the confocal laser at high intensity for 100 ms. The OFF response was recorded by turning off the lasers for 100 ms. The stimulations are contrast matched. Please see Methods for details. (**b**) Dark pulse-induced OFF responses were eliminated in the glu-lOLP by RNAi knock-downs of mAchR-B and Gαo. Average lOLP^glu^-Gal4>RCaMP traces are shown. The grey dashed line indicates the 100 ms dark pulse. The genotypes are as indicated. n = 7 in all groups. **(c)** RNAi knockdown of mAchR-B or Gαo demonstrates significant reductions in the amplitude of the calcium responses. Quantification of changed intensity (ΔF/F) is shown. Control, n = 8; mAchR-B^RNAi^, n = 6; Gαo^RNAi^, n = 7.

**Supplementary Figure 6.**
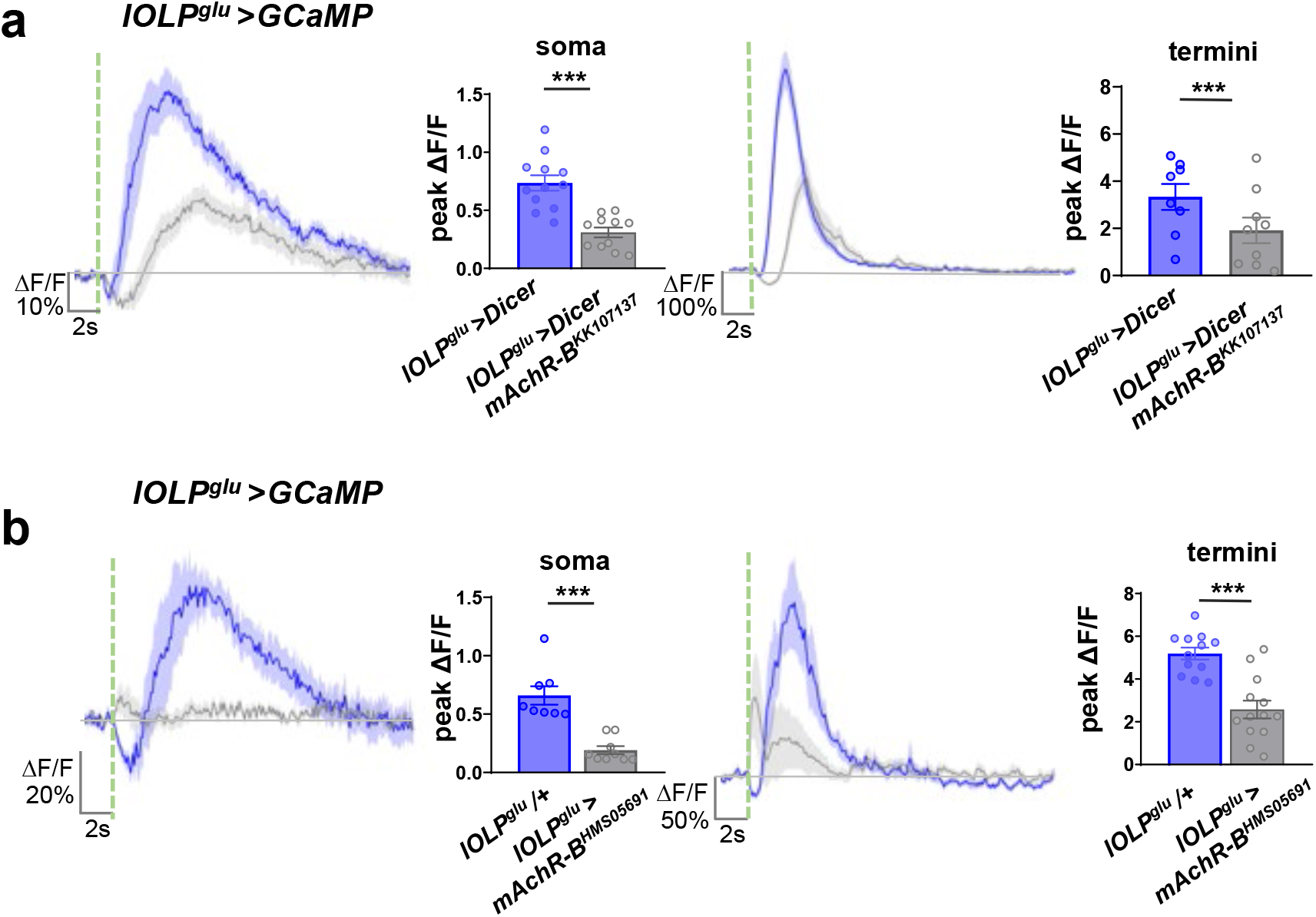
Knocking-down of mAchR-B consistently dampens the light-induced calcium responses in glu-lOLP. GCaMP signals were recorded in the soma (left) and termini (right) of glu-lOLPs, both of which showed significant decreases in amplitudes with the mAchR-B knock-down. Two transgenic RNAi lines, **(a)** mAchR-B^KK107137^ and **(b)** mAchR-B^HMS05691^, generated responses with different kinetics. Average traces and the quantification of peak changed intensity (ΔF/F) is shown. The green dashed line indicates the 100 ms light pulse. lOLP^glu^>Dicer, n = 12; lOLP^glu^>Dicer, mAchR-B^KK107137^ = 11; lOLPg^lu^/+, n = 8; lOLPg^lu^>mAchR-B^HMS05691^, n = 9.

**Supplementary Figure 7.**
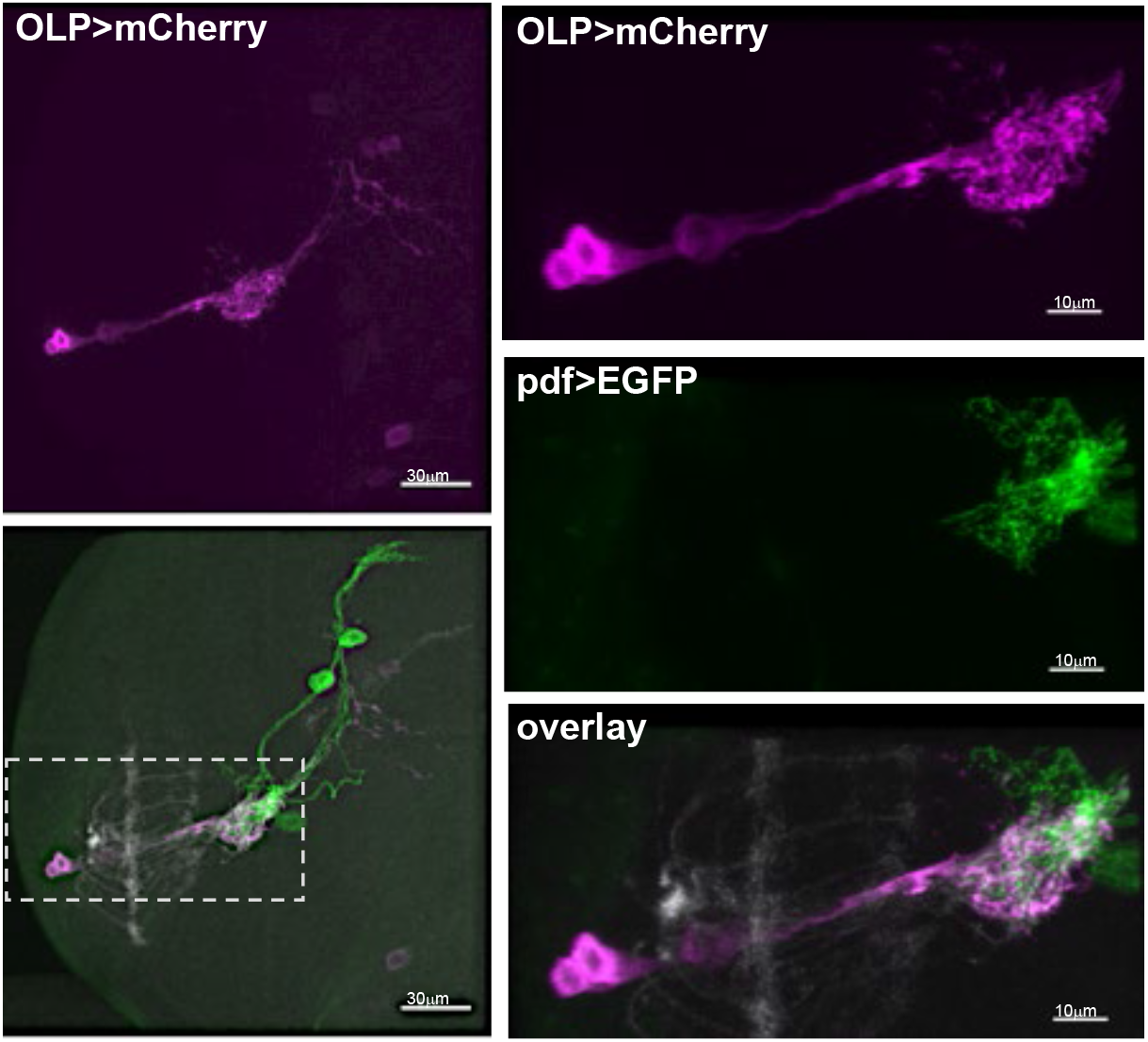
LNvs are visual projection neurons receiving synaptic input from PRs and OLPs. Representative confocal images illustrating the anatomical connections among PR axon terminals stained by 24B10 antibody (grey), the projections of OLPs (OLP-LexA>LexAop-mCherry, magenta) and dendritic arbors of LNvs (pdf>CD8::GFP, green) in the larval LON.

**Supplementary Figure 8.**
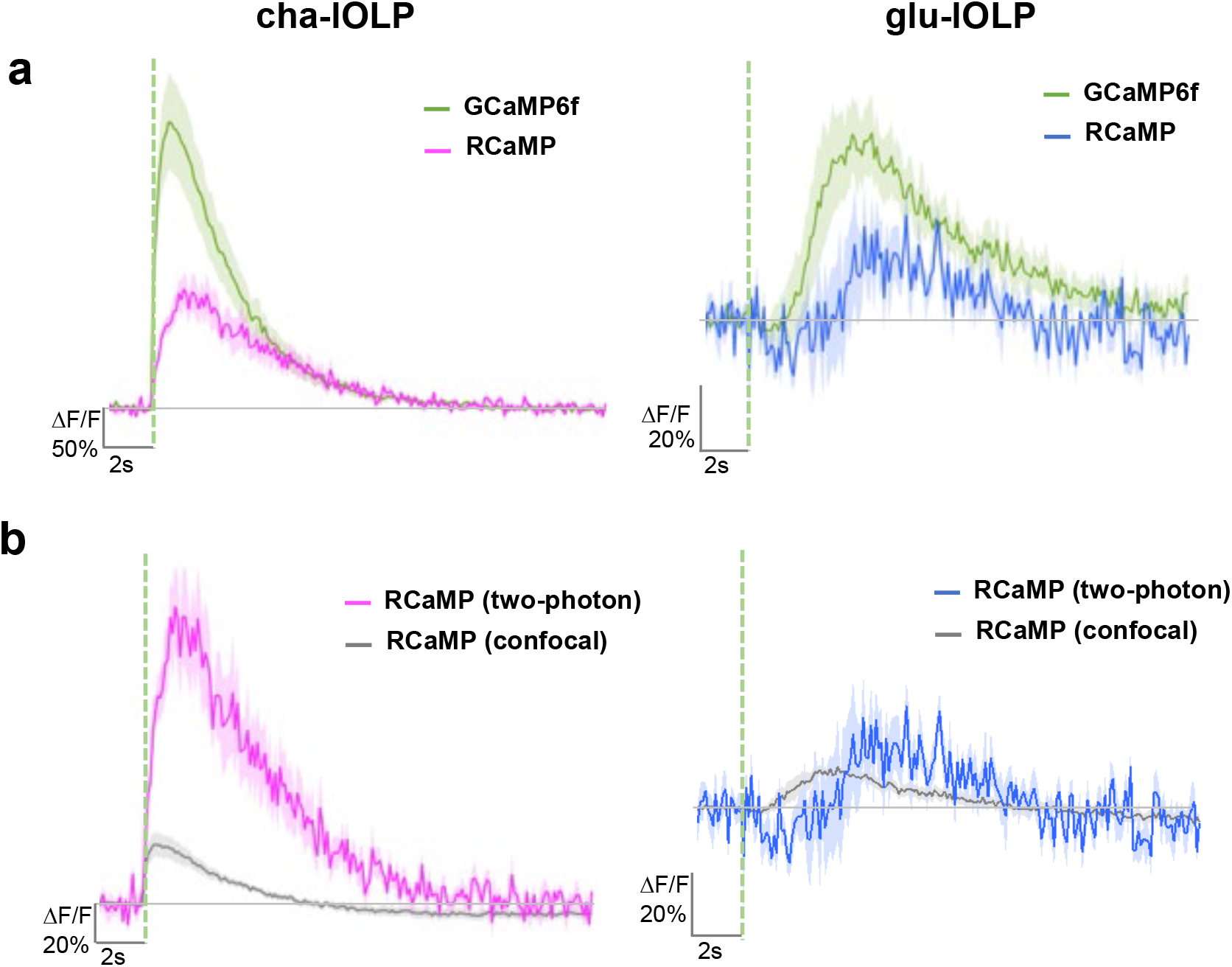
Comparison between calcium imaging results obtained using RCaMP vs. GCaMP6f and confocal vs. two-photon. **(a)** Calcium responses of the cha-lOLP (pink) and glu-lOLP (blue) were recorded using RCaMP or GCaMP6f with a two-photon laser tuned to 1040 nm or 920 nm, respectively. Although waveforms of the calcium transients are similar, the GCaMP recordings (green traces) showed responses with higher amplitude compared to the RCaMP recordings. **(b)** RCaMP recordings imaged using a two-photon laser tuned to 1040 nm produced calcium transients with higher amplitudes than the ones imaged using a confocal laser tuned to 561 nm (grey traces). The average traces are shown. Shaded areas represent SEM. 100 ms light pulses were delivered using the 561 nm laser and are indicated by the dashed lines.

